# Optimization and evaluation of complementary degrader discovery assays for application in screening

**DOI:** 10.1101/2025.03.06.641606

**Authors:** Johanna Huchting, Arjen Weller, Moyra Schweizer, Mona Brandt, Jan Heering, Maria Kuzikov, Markus Wolf, Jeanette Reinshagen, Markus A. Queisser, Philip Gribbon, Andrea Zaliani, Ole Pless, Aimo Kannt

**Author notes:** Corresponding author: Johanna Huchting.

## Abstract

Targeted protein degradation (TPD) mediated by molecular glues is an innovative pharmaceutical paradigm. By binding to and modulating the surface of an E3-ligase component, molecular glue degraders can facilitate the recruitment of a specific target protein (or vice versa) and, ultimately, invoke target degradation. This mode of action results in specific challenges for the development of rational discovery strategies, and complex hit validation workflows may be required to reliably eliminate compounds that elicit non-specific effects. With the aim to guide screening efforts, we optimized two orthogonal cell-based, target-centric assays for degrader discovery: (1) a time-resolved FRET assay directly quantifying levels of target protein and its degradation (signal inhibition), and (2) an assay coupling TPD to cell growth (signal rescue). To enable a deeper understanding of the individual assay’s strengths and limitations, we compared their statistical performance as well as respective hit populations by screening a specifically designed collection of about 1000 compounds containing well annotated reference compounds and known frequent hitters. We found that the signal rescue format reliably and specifically captured active target degraders while it efficiently filtered out interfering or frequent hitter compounds. Importantly, this format achieved to retrieve lower potency hits, which might be desirable in order to confidently include as many diverse chemical starting points as possible at the start of a drug discovery project.

## Introduction

Traditionally, small-molecule therapeutics subsume modulators of protein function such as inhibitors or activators. An increasingly important class of small molecules induces proximity between biological macromolecular entities, one being the biological target (e.g., a disease-modifying protein) and the other a biological effector. Molecular degraders, the most prominent type of proximity-based modalities, involve E3 ubiquitin ligases as biological effectors. They thus induce target protein (poly)ubiquitylation and subsequent proteasomal degradation, thereby depleting the protein of interest. Typically, the molecular degrader is retained and can thus induce degradation of the next target protein molecule, resulting in a catalytic mode of action.^1^ While some molecular degraders are already in clinical use, with the molecular glue degrader Lenalidomide (Revlimid®) as most prominent example, they have so far only retrospectively been shown to act via targeted protein degradation (TPD). Prospectively developed, advanced clinical candidates in this field largely belong to the heterobifunctional group of molecular degraders, such as PROteolysis-TArgeting Chimeras (PROTACs), that connect binders for target and E3-ligase via a linker.^2^ Molecular glue degraders (MGDs) can reach proteins that lack classical ligandable allosteric or orthosteric pockets, such as transcription factors, and are similar to conventional small molecule drugs in terms of classical drug-likeness. Systematic strategies for the prospective MGD discovery, however, are only beginning to emerge and rational design principles are not fully understood.^3–5^

When screening for molecular degraders with target protein abundance as primary readout, the presence of a target degrader results in a lower signal compared to vehicle control (signal inhibition), as would also compounds exhibiting rather unspecific effects such as cytotoxic compounds, translation inhibitors, etc. Consequently, stringent and extensive hit deconvolution cascades are needed to identify bona fide degraders from all hits of a screening library, and especially low-potency MGDs may be missed.^6^ In contrast to classical affinity-based modalities that follow an occupancy-driven mechanism, direct biophysical hit validation for catalytically acting degraders may be restricted.

Functional degrader validation generally includes recovery of target protein by proteasome inhibition or pharmacological inhibition of NEDDylation.^6^ However, in recent work Schwalm *et al*. have clearly demonstrated confounding effects of these strategies.^7^ Moreover, Vetma *et al*. have recently shown that cytotoxic compounds can disguise as targeted degraders even through profiling efforts, especially in the context of short-lived proteins.^8^

Our study compares target-centric, orthogonal assay formats for degrader discovery. It incorporates (1) direct measurement of target protein abundance (signal inhibition) as well as (2) an approach that couples TPD to a positive readout, specifically, cell growth recovery (signal rescue). The latter format was first introduced to TPD by Koduri *et al*. and employs exogenous expression of a target-suicide kinase fusion protein, a strategy borrowed from cell fate control gene therapy.^9^ We rigorously address challenges in assay sensitivity and robustness and provide detailed guidance on assay implementation. With the aim to understand the individual strengths and limitations of the different formats, we provide an in-depth analysis of screening results making use of a specifically designed validation library. This library leverages prior knowledge of the compounds’ biological effects as well as (predicted) compound promiscuity,^10^ i.e., frequently observed activities in a range of biological contexts. Such promiscuity may result from poly-pharmacology of a compound or pharmacological activity at central nodes in cellular pathways leading to more generalized downstream effects (e.g., translation inhibitors).^11^ Hence, our setup rigorously challenges the assay formats in terms of specificity and enables a concise interpretation of screening results. The findings from our study may guide TPD researchers during set up and rigorous vetting of assays for molecular degrader discovery as well as provide orientation for designing an efficient cellular screening architecture for the detection of molecular degraders.

## Results and Discussion

### Direct end-point measurement of target protein level

We implemented a homogenous time-resolved FRET (HTRF) sandwich immunoassay where, after treatment with a potential degrader, cells are lysed and a FRET-labeled antibody pair is added to detect separate epitopes on the same target protein. The assay does not require any washing after cell lysis and the read FRET signal is directly proportional to the amount of target protein in the sample. As target, we selected the transcription factor IKZF1 (IKAROS family zinc finger 1). IKZF1 is targeted by numerous thoroughly characterized MGDs that utilize the E3 ligase component cereblon (CRBN) and are therefore being referred to as cereblon-E3-ligase-modulating drugs (CELMoDs).^12–19,20(review)^ A central aim of this study was a direct comparison of this signal inhibition assay to an orthogonal, signal rescue-type format, and hence, the choice of cell line had to consider the needs of both assays. For the signal rescue format, the assay principle relies on the expression of a target-suicide kinase fusion protein; consequently, we adapted and optimized both assay formats using the same, genetically modified cell line. Following assay miniaturization to a 384-well format and automation, and with our focus set on achieving a robust setup suitable for screening, we then aimed to maximize the assay window (directly depending on maximal degradation%, D_max_) and minimize variation. For this, a 20 h treatment with pomalidomide seems best suited (Figure 1A), while for other target/degrader-pairs, 6 h treatment may already suffice (as we found for RBM39/indisulam, data not shown). Relative IKZF1 protein levels measured after 20 h treatment with diverse CELMODs reflected well the published activities of these compounds (Figure 1B).

**Figure 1:**
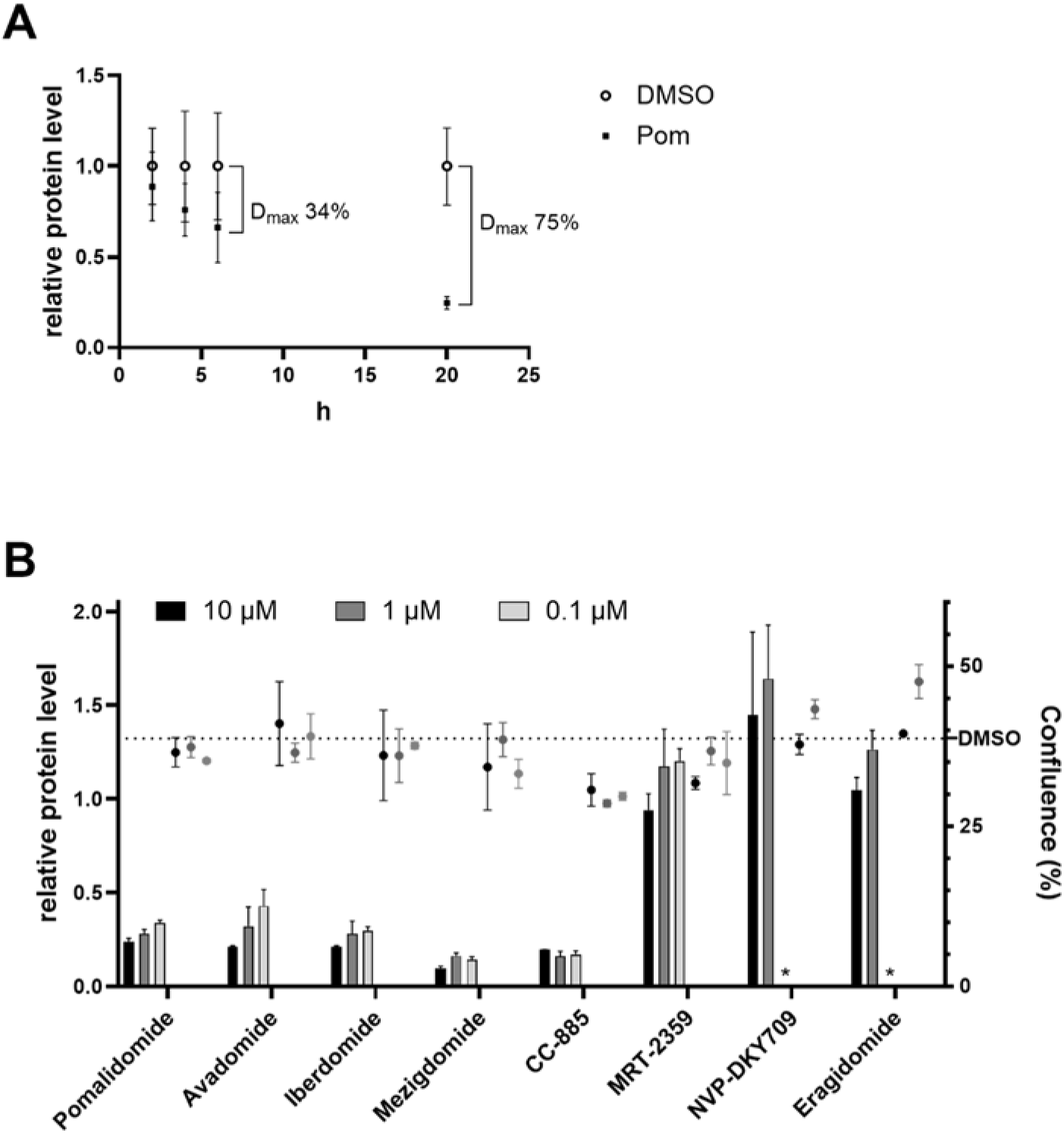
Relative IKZF1 protein level measured by HTRF. **(A)** Multiple endpoint study: Maximal degradation reaches 11, 24, 34 or 75% after 2, 4, 6 or 20 h treatment with 10 µM pomalidomide. **(B)** Compound profiling: Relative protein level after 20 h treatment with different CELMoDs recapitulates published compound activities (left y-axis, measured by TR-FRET and normalized to DMSO vehicle). Confluence (%) was measured in parallel to signal potential toxic effects, as may be the case for CC-885 (right y-axis, measured via brightfield microscopy). Data are means of N=3±SD. *Condition not included

Aiming to understand the assays’ strength and limitations for screening medium-sized chemical libraries, we designed a validation library (Figure 2). Briefly, this library of 941 small molecules was assembled from (i) annotated compounds (∼1/3 of the set) that have extensively been studied both in biochemical as well as functional assays, with a well-understood bioactivity and mode of action including known molecular degraders of diverse proteins engaging different E3 ligases (both bifunctionals and glue degraders; “bioactives” in Figure 2) and (ii) frequent hitters (FHs), i.e., compounds with a high likelihood of appearing active in various biological assays without following the sought mechanism (as predicted by the Hit Dexter machine learning model;^10^ ∼1/3 of the set). Finally, this validation library included (iii) a subset of structurally blinded proprietary compounds (∼1/3 of the set) that consisted of small molecules with well-understood, specific bioactivities. This latter part of the library was not explicitly enriched in targeted molecular degraders but may have included CELMoDs at low proportion, hence overall reflecting the makeup of the disclosed bioactives subset and serving to test our setup (Table S1; disclosed compounds SMILES in supporting information xlsx).

**Figure 2:**
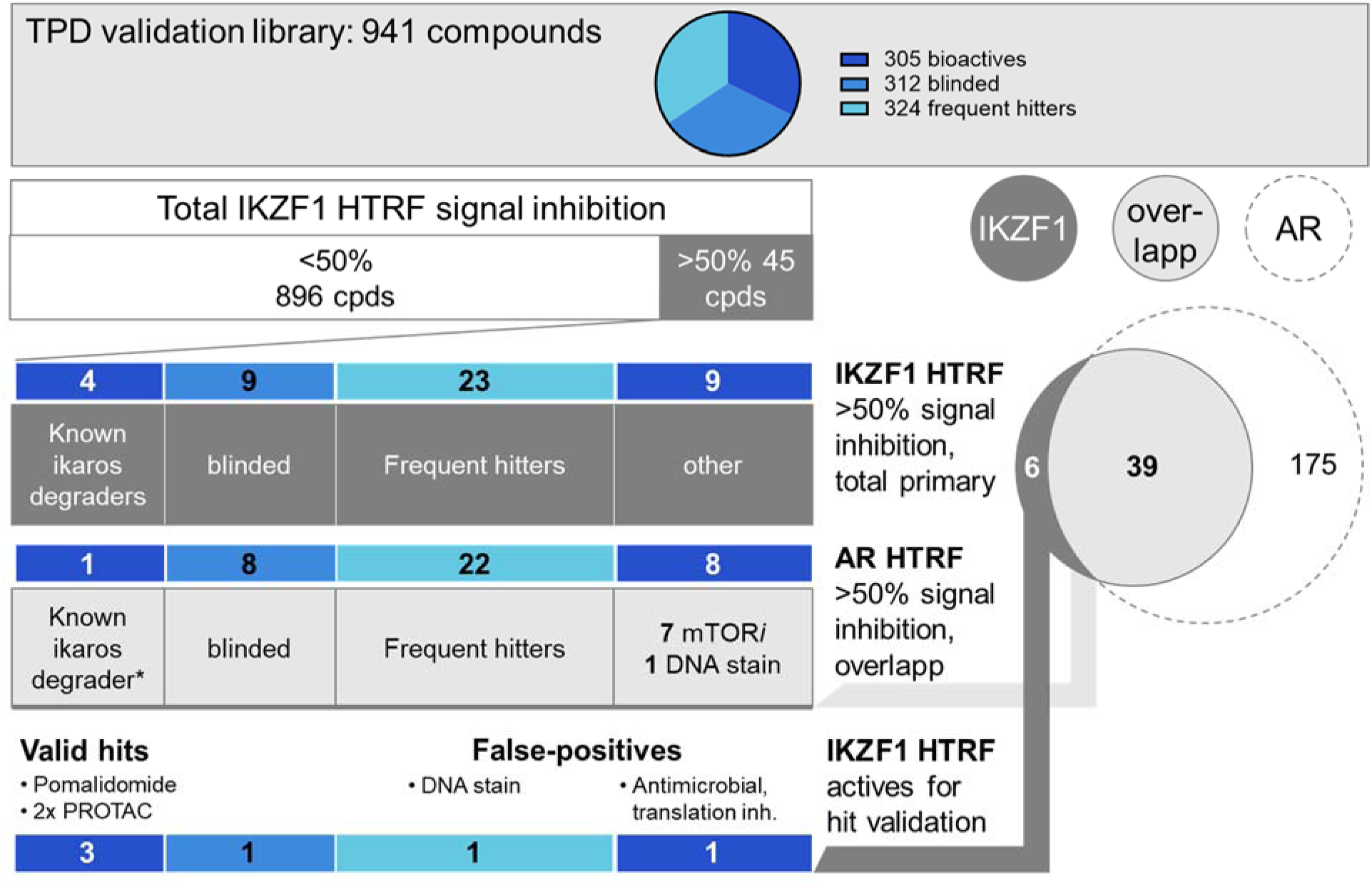
TPD validation library subsets and analysis of primary hits from the IKZF1 HTRF screen. Primary hits are defined as inhibiting the signal by at least 50% after signal normalization (0%inhibition – vehicle, 100%inhibition – pomalidomide (IKZF1) or ARCC-4 (AR)). AR=androgen receptor

In screening mode, the assay achieved acceptable statistical consistency^21^ and 45 primary hits were identified (Figure 2; Z’-values ranged from 0.5 to 0.6). Among these primary hits were pomalidomide as well as three PROTACs that incorporate pomalidomide-derived E3 binders and thus induce degradation of IKZF1 as off-target. After removal of 9 blinded compounds, the remaining 32 primary hits contained 23 FHs according to the cell-based prediction model^10^. These included inhibitors of topoisomerase, protein synthesis, or ATPases and fluorescent compounds/DNA intercalators. Of the remaining 9 hits the majority were inhibitors of mTOR that were not flagged by the FH prediction model.

Assuming that the same assay principle, screening workflow and library applied to an unrelated target may serve as a counter strategy to invalidate hits, we analyzed the overlap of primary hits from the IKZF1 HTRF screen with a similarly executed HTRF screen that measured endogenous androgen receptor (AR) levels. This analysis identified 39 shared primary hits whereof the majority were predicted FHs^10^ or known to inhibit mTOR (Figure 2). Hence, while the primary hit rate was at 4.8%, the counter assay flagged almost 90% of these hits as unspecific, leaving six for downstream validation. From these, three compounds were confirmed IKZF1 degraders while the remaining two false-positives included a DNA stain and a translation inhibitor (one compound was blinded). However, this validation strategy would not necessarily have worked the other way round, using the overlap with the IKZF1 screen to remove unspecific compounds from the AR screen, as in the latter case the primary hit rate was higher (23%).

In summary, we implemented a homogenous sandwich immunoassay that achieves specific detection and reliable quantification of target protein directly from cell lysate, thereby enabling degrader discovery in a screenable format without the need for tagged proteins. While this assay captured known target degraders from the validation library, primary hits were enriched in FHs and included compounds with generalized effects such as translation inhibition/cell cycle arrest. These false-positives were largely invalidated through a similarly executed screen on a different target.

### Degrader-induced recovery of cell growth

An alternative approach to eliminate compounds with generalized effects such as translation inhibition from primary hits in degrader screening is the signal rescue format first introduced by Koduri *et al*.^9^ This assay couples TPD to a positive readout, specifically restoration of cell growth.^22–25^ The assay principle relies on fast-growing cells that are stably transduced to bicistronically co-express i) the target protein fused to a suicide kinase (dCK*) and ii) eGFP, serving as an expression marker both during cell line generation and degrader screening. The suicide substrate 5-bromovinyl uridine (BVdU) selectively inhibits cell growth in dCK*-positive cells while dCK*-negative cells remain unaffected. In cells expressing dCK*-target fusion protein, growth is recovered by a molecular degrader of the target protein due to co-degradation of dCK* (Figure 3, top). This way, compounds with described generalized effects would not be identified as hits. A detailed description of the assay principle is provided in supporting information (pdf).

**Figure 3:**
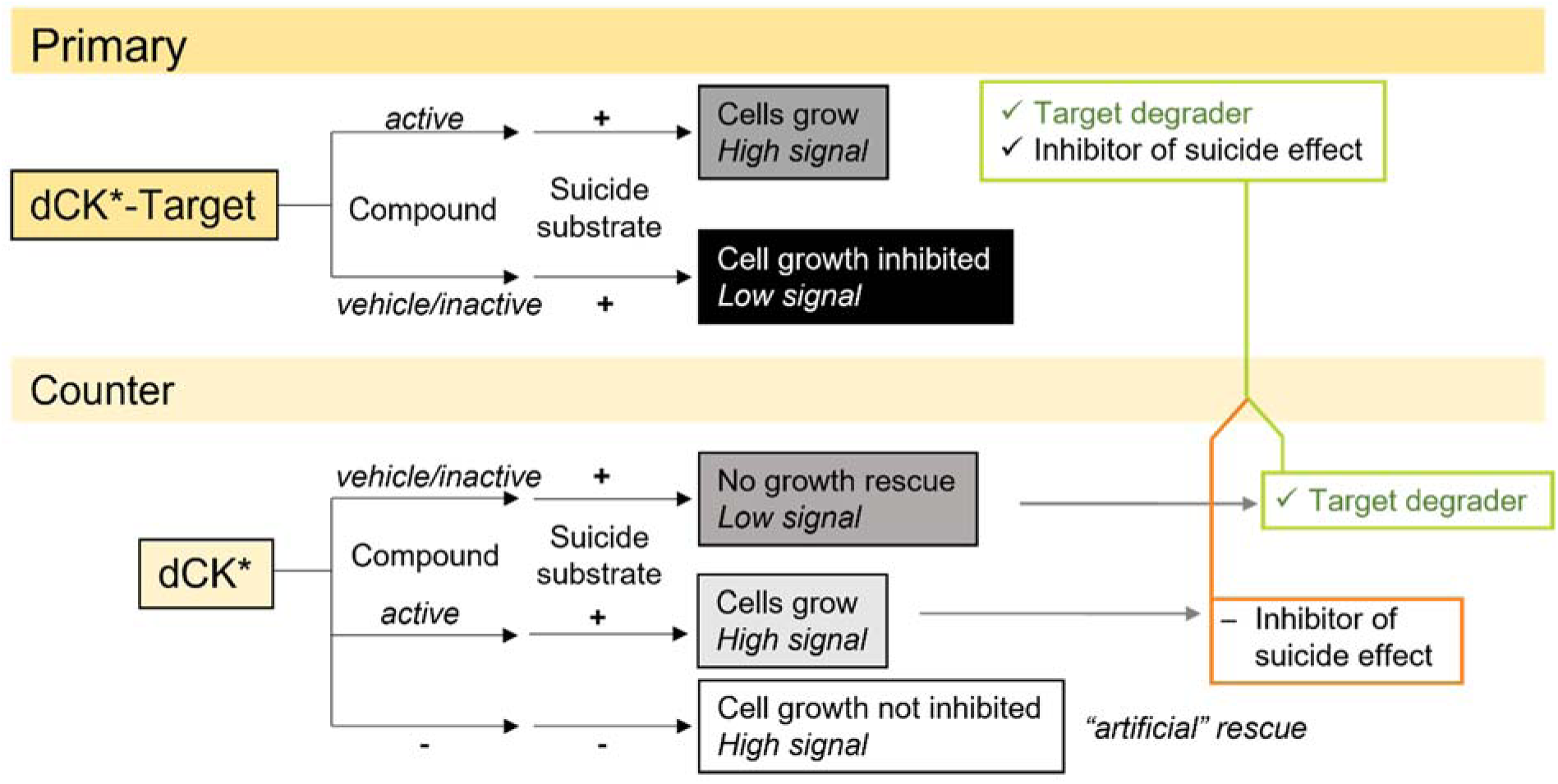
Assay principle of the growth recovery assay. The primary readout of this format measures cell growth at five days after treatment initiation and hence, compounds perturbing essential cellular functions will not be among primary hits. The counter assay invalidates primary hits that target suicide kinase-related components.

As this assay in its originally described version in our hands did not reliably achieve sufficient assay window and robustness for single-point screening, we extensively optimized the protocol while maintaining its rational basis. Briefly, GFP-based object counting in our hands underestimated the actual cell number present in the well. This underestimation was especially pronounced at higher cell densities and consequently, the high control signal was artificially lowered. To improve object separation during image analysis while keeping acquisition times short, we switched the assessment of cell growth to Hoechst nuclear staining-based object counting, which more faithfully reflected cell count and thus increased the assay window by over three-fold (Figure 4A).

**Figure 4:**
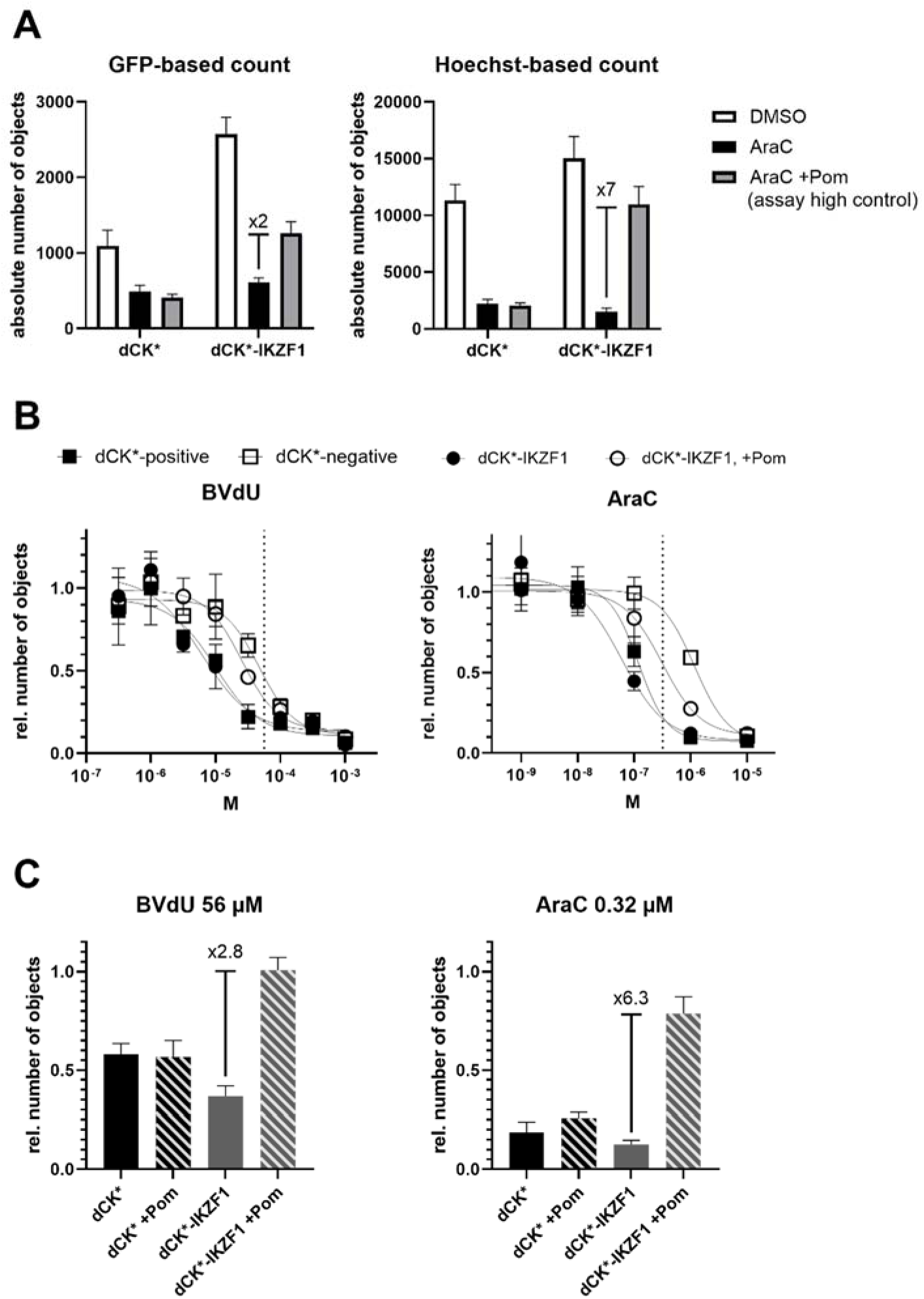
Optimization of the assay window. **(A)** Comparison of GFP-based *versus* Hoechst-based object counting. Absolute objects are retrieved from images of whole wells in 384-format before (GFP) or after live-cell nuclear staining (Hoechst 33342) (N=8±SD). **(B)** Concentration-dependent BVdU or AraC-induced suicide effect in dCK*-positive (′) versus -negative cells (≤) and in dCK*-IKZF1 cells pre-treated with DMSO (●) versus pomalidomide (■) (N=3±SD). Dotted lines indicate concentrations chosen for the screening assay. **(C)** Comparison of suicide substrate effect in screening layout. Pomalidomide pre-treatment shows a significant rescue effect in dCK*-IKZF1 but not in dCK* cells under both suicide substrate conditions, yielding the largest assay window for 0.32 µM AraC. In B and C, objects represent nuclei and are retrieved via live-cell Hoechst-staining and image-based counting from whole wells in 384-well plate format (N=). Note that in concentration-response experiments, both BVdU and AraC appear to have increased potency due to the difference in final DMSO (B:1.05%, C:0.15%).

Furthermore, we enhanced BVdU potency by co-treating cells with TAS-114, a dual dUTPase/ dihydropyrimidine dehydrogenase inhibitor, and achieved an almost 4-fold assay window (Figure S1).^26^ Still, aiming for an effective suicide substrate, we finally substituted BVdU with the nucleoside analogue cytarabine (AraC), an antineoplastic anti-metabolite.^27^ In concentration-response experiments, we found that AraC was more potent than BVdU (2-log difference in IC_50_-values). Importantly, AraC achieved adequate selectivity, probably due to the overexpression of constitutively active kinase dCK* rather than differences in substrate preference for dCK* versus cellular dCK (Figure 4B).^25^ Finally, in screening layout, this setup achieved an assay window of 6-fold under 0.32 µM AraC-treatment, which we could not reach with BVdU (Figure 4C).

Next, we assessed the growth recovery assay performance in degrader profiling (Figure 5). As anticipated, the results from the recovery assay presented a nearly opposite scenario compared to the signal inhibition configuration with IKZF1 degraders giving high signal. Chemically related compounds that are known to leave IKZF1 levels unaffected yielded low signal and, interestingly, compounds targeting proteins that are essential for cell growth even scored below baseline (Figure 5B). This effect was more pronounced with longer compound pre-treatment and may assist in interpretation of screening results (Figure 5C).

**Figure 5:**
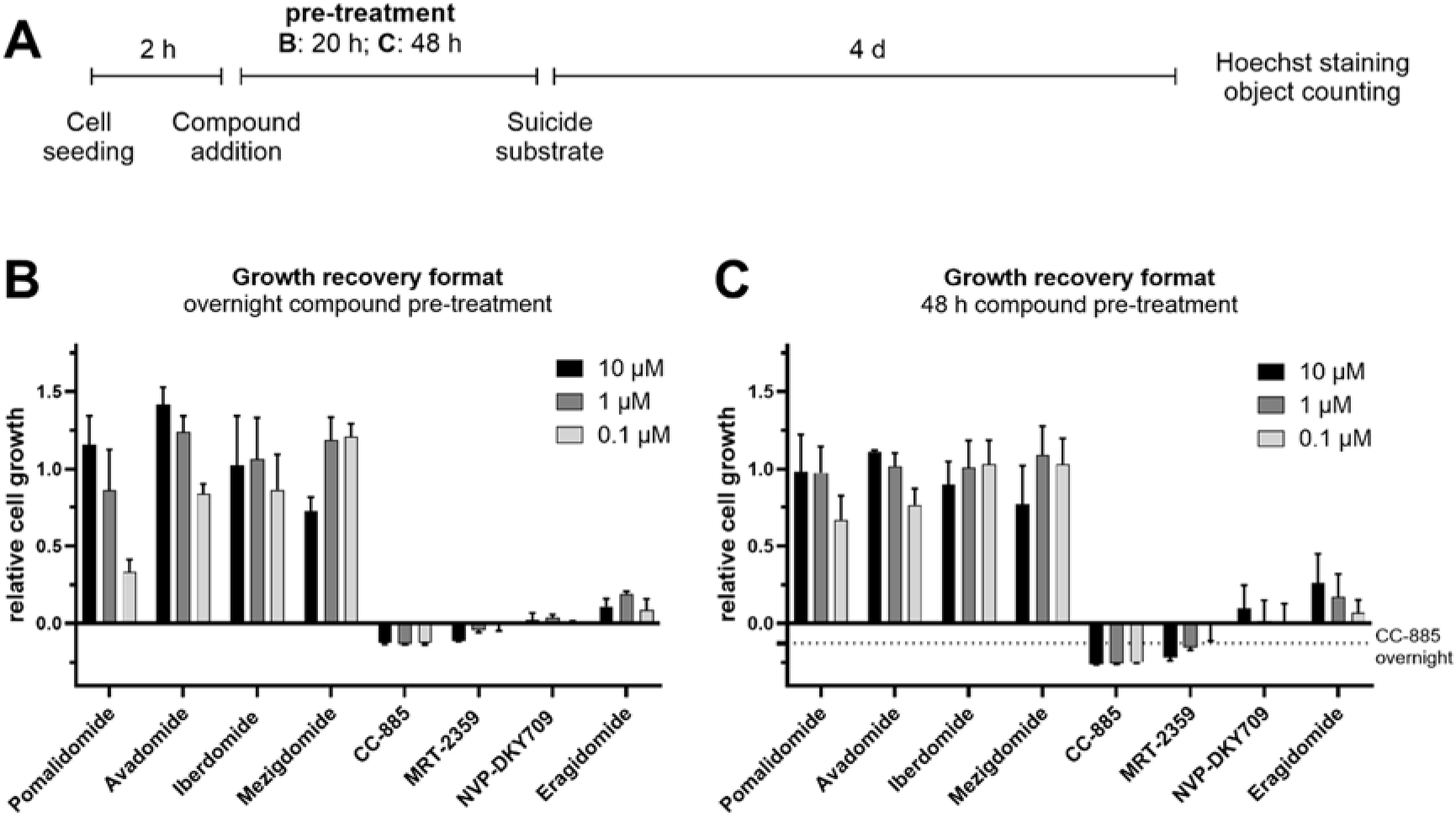
Growth recovery of dCK*-IKZF1-modified cells by different CELMoDs. **(A)** Consecutive steps and timeline of the growth recovery assay. **(B)** Relative cell growth determined via Hoechst-based object counting after 20 h compound pre-treatment followed by four days of suicide substrate (AraC)-treatment (normalized to pomalidomide-treated control wells as full growth recovery and vehicle-treated wells as negative control). **(C)** Variation of A where compound pre-treatment time was increased to 48 h.

When applied for screening the same TPD validation library as described for the FRET-based assay, the growth recovery format performed robustly as well (Z’-values of 0.4-0.5).^21^ While for the primary assay in dCK*-IKZF1-modified cells, pomalidomide served as on-target growth recovery control, there was no such control available for the counter assay in dCK*-modified cells. The original reported protocol used dipyridamole, a nucleoside transport inhibitor;^9^ however, this approach was incompatible with our readout as it completely prevented Hoechst staining and, in our hands, increased GFP intensity compared to untreated control, an observation we did not follow up on. Hence, for the counter assay, non-AraC-treated wells served as simulated growth recovery control. Due to these differences in normalization, thresholds for hit detection varied slightly between primary and counter screen. The resulting 7 hits included pomalidomide from within the library and library-PROTACs that incorporate pomalidomide-like E3-binders, hence capturing an off-target effect of CRBN-engaging PROTACs (Figure 6A, green stars). As expected, PROTACs engaging other E3-ligases and, importantly, those targeting CRBN through advanced thalidomide 5-fluoride-derived binders (e.g., ARV-110) were not identified as hits. This was confirmed in concentration-response experiments (data not shown).

**Figure 6:**
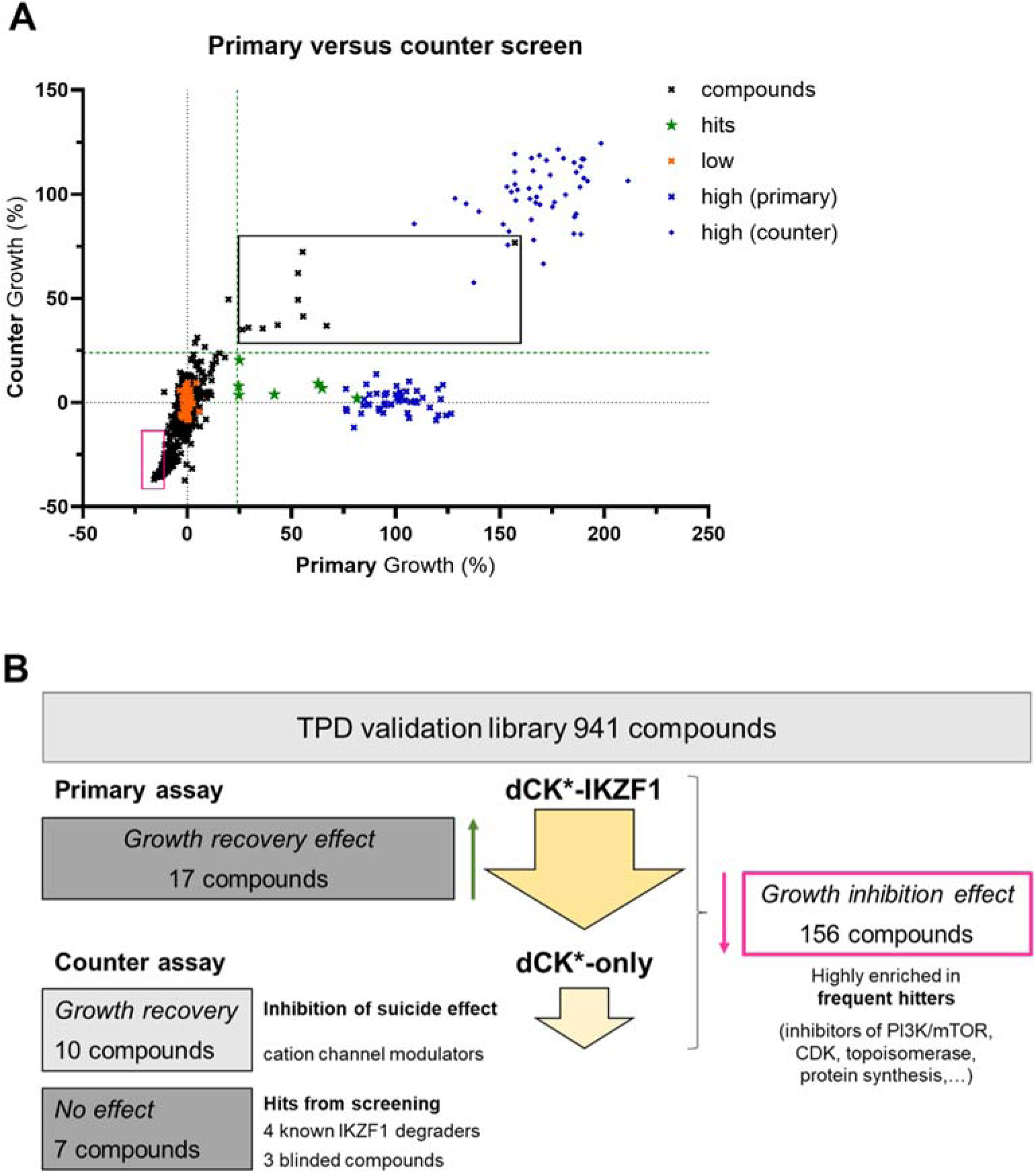
TPD validation library screening in growth recovery assay. **(A)** Relative growth recovery by library compounds or controls in primary versus counter assay. Primary assay: dCK*-IKZF1-modified cells, 100% growth recovery is achieved by 10 µM pomalidomide (x-axis); counter assay: dCK*-modified cells, “no suicide”-control as 100% growth (simulated recovery, y-axis). Compounds from the TPD validation library were screened at 5 µM. Hits from the primary screen that were not invalidated by the counter screen are highlighted as green stars while those that were also active in the counter screen are within the black rectangle. Compounds within the section in magenta show a marked growth inhibitory effect in both, primary and counter screen. **(B)** Schematic overview of the compound-focused analysis of results from the growth recovery primary versus counter screen.

To gain a detailed understanding of the strengths and limitations also of this format, we analyzed those hits from the primary screen that were invalidated by the counter assay (total of 10 compounds; Figure 6B). Among these compounds, cation channel modulators were present (lidoflazine, FPL64176, SR1078, A-784168). In line with the compounds’ signal rescue effects in dCK*-only cells, the variety of agonist/antagonist modulation and the structural diversity found in these compounds suggests higher probability of target-unrelated cellular effects (ultimately leading to an inhibition of the suicide effect) rather than selective modulation of IKZF1-related cellular mechanisms. Accordingly, the majority of these false-positives were also active when screening a different target in this format (not shown).

Importantly, a significant proportion of the library compounds elicited substantial growth inhibitory effects on top of the AraC-induced inhibition, both in the primary and counter assay (Figure 6, magenta). This effect had also been noted for some of the CELMoD’s described above, most prominently CC-885. With the aim to potentially exploit this “multiplexing” of growth inhibitory and recovery effects, we performed a detailed analysis of the inhibitory compounds (Figure 6B and Table S2). Among the 156 compounds that yielded below -11% growth, 28 structures were blinded (i.e., ∼10% of the blinded subset elicit this inhibitory effect). Out of the remaining 128 non-blinded compounds, only 25 were from the disclosed bioactives subset, confirming that ∼10% of non-promiscuous bioactives elicit an inhibitory effect in this assay. These included assay principle-related inhibitors (cyclocytidine: a prodrug form of AraC, probably enhancing the AraC-effect by increasing its final concentration; decitabine: a DNA-methyltransferase inhibitor that requires activation by dCK), and inhibitors of central cellular nodes (topoisomerase, PI3K/mTOR and CDK). As expected, from the frequent hitter^10^ subset of the library we find growth inhibitory compounds at an increased rate of ∼30% (103 compounds). These included members from the anthracycline class or polycyclic quinoids (e.g., epirubicin, mitoxanthrone), other topoisomerase inhibitors as well as inhibitors of protein synthesis.

### Orthogonal degrader assays in a screening cascade

Finally, we performed another small-scale screen of a different compound set with a similar size (1001 small molecules, SMILES in supporting information xlsx). Molecules were selected based on structural features known from the few well-established molecular glue degraders (e.g., aryl sulfonamides, phthalimides, pyridyl thiazole amines, triazino indoles etc.). As a first, strong and cost-efficient filter, we employed the growth recovery assay (primary and counter), to then use the FRET-based format for hit profiling. The growth recovery screening in dCK*-IKZF1 versus dCK*-only cells resulted in 9 hits (0.9% hit rate; Figure 7B, green stars). While the majority of compounds in the set do not have related bioassay annotation data, some were known thalidomide analogues with reported activities in relevant assays. For example, 4- and 5-hydroxythalidomide have been reported inactive towards IKZF1- degradation in a structure-activity relationship study while still binding CRBN^19^ and in accordance with reports, these compounds were not among hits in the screen indicating the suitability of our assay as a source of TPD mediated primary hits.

**Figure 7:**
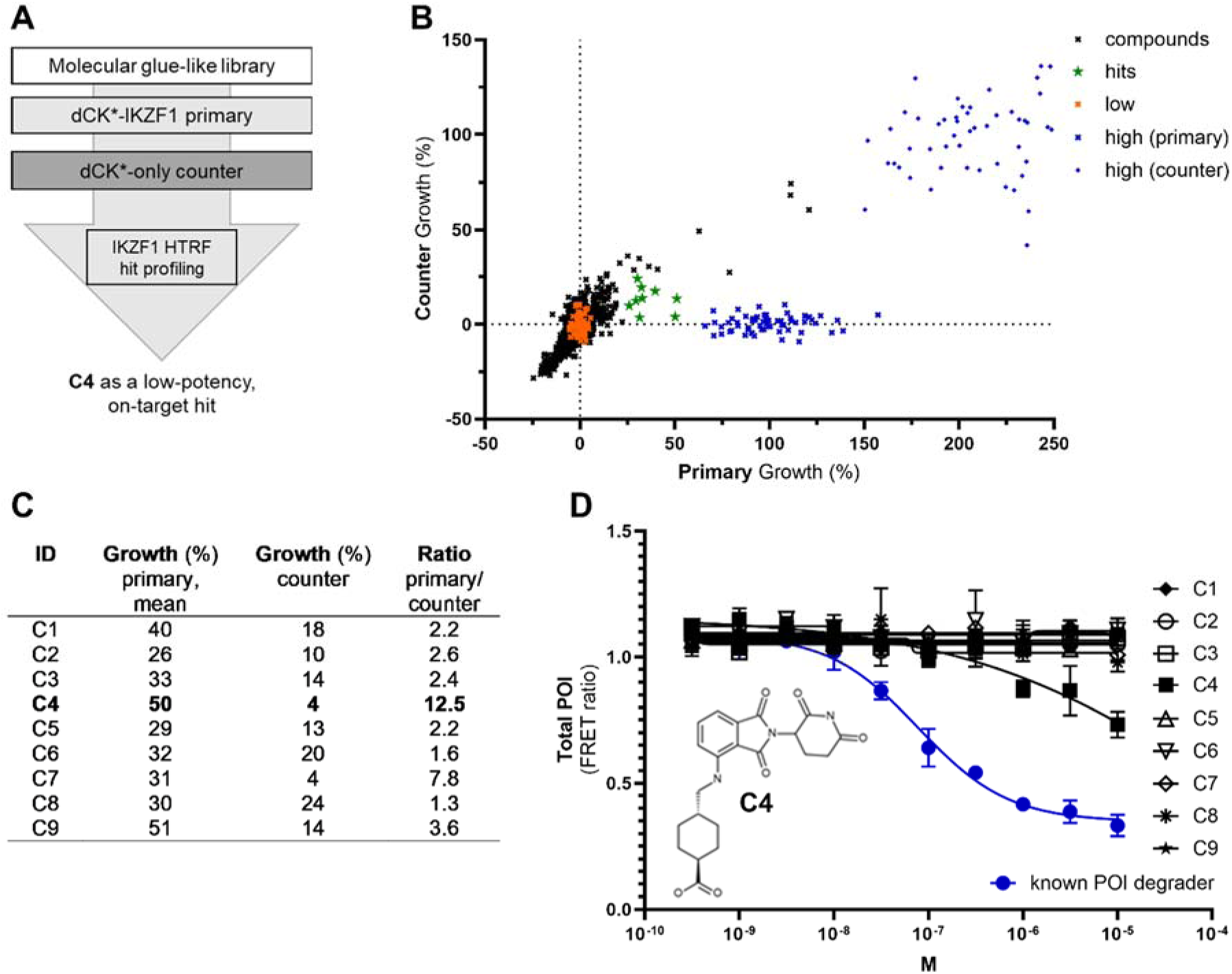
Proof-of-concept study. **(A)** Overview of the screening flow. **(B)** Relative growth recovery by compounds or controls in primary (dCK*-IKZF1; x-axis) versus counter (dCK*-only)-setup (y-axis). Library compounds (1001 small molecules) were screened at 5 µM. 9 Hits were identified (green stars). **(C)** Table showing growth (%)-values from primary (N=2) and counter as well as the calculated selectivity ratio for the 9 hit compounds. **(D)** Profiling of the 9 hits with the FRET-based assay in triplicate 10-point half-log concentration-response and in direct comparison with pomalidomide as known IKZF1 degrader; molecular structure of C4.

Next, all 9 hits were profiled in triplicate 10-point half-log concentration-response curves using the FRET-based assay (Figure 7D). Importantly, general inhibitory effects were already excluded as all 9 compounds had passed the signal rescue filters, where, as established above, growth recovery cannot be achieved with compounds exhibiting, e.g., general inhibition of protein synthesis. False-positives from the primary screen may include compounds that interfere with the expression of the integrated exogenous gene more specifically, as well as those leading to degradation of the fusion protein not via IKZF1 but via dCK*. These compounds, however, would also be active in the dCK*-only counter screen and were thus already invalidated. The most potent and selective of the 9 hit compounds (compound C4 in Figure 7) showed concentration-dependent inhibition of FRET ratio, indicating a likewise decrease in target protein levels. This validated hit compound C4 incorporates the 4-amino-substituted phthalimide scaffold of pomalidomide, yet it is less potent; in fact, C4 might have been overlooked in a direct FRET-based screen. It is generally desired to efficiently identify even low-potency but “on-mechanism” hits since after screening (in hit-to-lead development), medicinal chemistry strategies can increase potency of otherwise favorable compounds (e.g., regarding “on-mechanism” feature, scaffold novelty). A scenario where primary screening provides less efficient hit filtering may not allow to set the hit threshold at levels that include these low-potency, “true positive” hits, simply because the high number of hits implies a high proportion of false-positives and necessitates extensive further hit validation. As a consequence, promising chemical starting points may be missed (false-negatives).

## Conclusions

Others have highlighted the significant challenges in designing and interpreting cascade experiments for identifying, triaging and validating hits from a degrader screening campaign, ultimately aiming to identify bona fide molecular degraders,^6,8^ and it appears highly desirable to incorporate high fidelity assays with interpretable outcomes early in these screening workflows. We have first rigorously addressed critical parameters for the implementation of degrader assays for screening. Secondly, our validation library was specifically designed to analyze hit populations arising from different TPD assays used for screening to ultimately enable faithful interpretation of screening results. This allowed the side-by-side evaluation of two complementary assay formats regarding performance and hit populations, revealing their individual strengths and limitations and thus guiding the design of a proof-of-concept study. Here, compound C4 was identified as a bona fide, although low-potency degrader of the transcription factor IKZF1. In conclusion, our results clearly illustrate that the cascade and filters applied are highly robust and sensitive to efficiently identify even low-potency on-target and on-mechanism chemistry. Further work including different targets and a larger screening set are needed to provide an estimation of the general applicability of this approach and, during hit validation, compound triaging may benefit from consideration of degradation kinetics.

## Methods

Suppliers and catalogue numbers for all commercial material used in this study are provided in supporting information (pdf).

### Cell lines and culture

HEK293FT cells were grown according to manufacturer’s specifications in complete medium consisting of Dulbecco’s Modified Eagle Medium (DMEM; high glucose) supplemented with 10% FBS, 6 mM L-glutamine, 1% penicillin/streptomycin, 1x MEM non-essential amino acids and 1 mM sodium pyruvate at 37 °C and 5% CO_2_. To maintain growing cultures of unmodified HEK293FT cells, complete medium was supplemented with 500 µg/mL G-418, and for growing cultures of stably transduced, HEK293FT- derived cell lines, 8 µg/mL Blasticidin was added on top of this. During assays, no selection antibiotics were present.

For details on generation of stably transduced cell lines see supporting information (pdf).

### Compounds used in this study

The small molecule library we used as a platform for initial screening (validation library composition in supporting information xlsx) is composed of 941 compounds, partly derived from in-house drug repurposing collections^28^, where a curated database is publicly available listing the compounds, indications, primary targets (where known) and mechanism of action.^29^ Additionally, known molecular degraders (Molport) and structurally blinded compounds from a proprietary, well-annotated Chemical Biology library were included. A third subset consisted of predicted frequent hitters (identified from in-house small molecule libraries via an ML prediction algorithm for frequent hitters, Hit Dexter 2;^10^cell-based assay nuisance compounds or potential degraders).

The small molecule library used to test our optimized assay setup is composed of 1001 compounds that were selected based on scaffold similarity to published molecular glue degraders (Specs). Scaffolds present in the set include aryl sulfonamides, pyridyl thiazole amines, triazino indoles, phthalimides. All known compounds structures are disclosed in supporting information xlsx.

Other small molecule compounds used in this study were commercially sourced (supporting information pdf).

### Homogenous sandwich immunoassay

#### General procedure

The assay was performed using the HTRF Human and Mouse Total IKZF1 Detection Kit (Revvity, 64IKZF1TPEG) according to the manual. Briefly, stably transduced dCK*-IKZF1-HEK293FT cells were seeded in 20 µL complete medium per well in 384-well plates (SpectraPlate-384, Revvity, 6007650) and allowed to adhere overnight at 37 °C and 5% CO2 before compound from DMSO stock or vehicle were added using an Echo acoustic dispenser (Labcyte). Plates were incubated overnight at 37 °C and 5% CO_2_. Next, the treatment medium was slowly aspirated from all wells using a Janus MDT liquid handling station equipped with 384-well robotic tips (Axygen, Corning, PK-384-R). Lysis buffer was dispensed manually (20 µL per well). Plates were briefly centrifuged and incubated with lid at room temperature for 30 min. The detection antibody mix (3 µL/well) was manually dispensed into detection plates (ProxiPlate 384-shallow well Plus, Revvity, 6008280). Next, using the Janus MDT station equipped as above, cell lysates were homogenized by carefully pipetting up and down and lysates transferred to the pre-filled detection plate (12 µL/ well). Plates were sealed and incubated overnight before TR-FRET signal was read on an EnVision multimode reader (Revvity; Flash lamp, 100 flashes; Excitation: 320 nm, bandwidth 75 nm; Emission: channel 1 620 nm, bandwidth 10 nm; channel 2 665 nm, bandwidth 7.5 nm; Delay 60 ms). FRET ratio was calculated as channel2/channel1 and background subtracted data were normalized to vehicle (DMSO, 0% inhibition) and positive control (pomalidomide, 100% inhibition). Wells for background did not receive cells but included complete medium, DMSO-treatment, equivalent work-up and antibody mix.

#### Concentration-response experiments

To evaluate the inhibitory effect of compound treatment on dCK*-IKZF1 protein level, quantified via the FRET ratio, 2000 dCK*-IKZF1 cells/well were seeded according to the general procedure and 10- point half-log serial dilutions starting from 10 µM (20 nL from 10 mM DMSO stock) were added the next day. Per plate at least 8 pomalidomide-treated (10 µM) and at least 8 DMSO-treated wells were included for normalization.

#### Screening

According to the general procedure, 2000 cells/well were seeded in columns 2-24, column 1 received complete medium. Library compounds were dispensed into single wells at 5 µM final concentration (10 nL from 10 mM DMSO stock) in plate columns 3-22. Columns 1 and 24 received DMSO (10 nL) and column 23 received 10 µM pomalidomide (10 nL from 20 mM DMSO stock).

### Growth recovery assay

#### General protocol

Stably modified HEK293FT cells were seeded in 20 µL complete medium at a density of 300 cells/well in 384-well imaging plates (PhenoPlate 384-well; Revvity 6057302) using a Multidrop Combi+ dispenser (ThermoFisher Scientific) equipped with a standard cassette (Cat.no. 24072670). At two hours after seeding, compounds or vehicle were added using an Echo acoustic dispenser (step 1). Plates were incubated overnight at 37 °C and 5% CO_2_ before suicide substrate was added via acoustic transfer (step 2). To allow for optimal cell growth during the following 4 days, 20 µL complete medium was added using a Multidrop Combi+ dispenser, and plates were again incubated at 37 °C and 5% CO_2_. At day 5 post cell seeding, 10 mg/mL Hoechst 33342 in dd water was diluted 1:1000 in PBS and all wells received 10 µL per well of this dilution using a Multidrop Combi+ dispenser (final Hoechst concentration 2 µg/mL). Plates were incubated for 45 min at 37 °C and 5% CO_2_ before being imaged (EnSight multimode plate reader, Revvity; inverted optical microscope with 4x magnification and sCMOS image sensor). Objects were detected in full wells (BLUE channel, excitation 385 nm, 30 ms, 100%) and counted (Kaleido 3.0, Object detection method C).

#### Concentration-response experiments: suicide substrate

To evaluate the potency of 5-bromovinyl-deoxyuridine (BVdU) and cytarabine (AraC), 10-point half-log dilutions (BVdU) or log-dilutions (AraC) starting from 0.1 M were prepared manually in DMSO. The assay was performed according to the general protocol described above using, in step 1, 10 mM pomalidomide (20 nL of 10 mM DMSO stock) as signal rescue control for dCK*-IKZF1 cells and vehicle (DMSO) for all other conditions. On the next day, 400 nL of suicide substrate dilutions were dispensed in quadruplicate (step 2) and the general protocol was followed (final concentration ranges: BVdU 1 mM - 30 nM, AraC 1 mM - 1 pM; final DMSO 1.05%). For all cell lines, normalization was based on object count averaged from at least 8 wells that received vehicle only (100% growth).

#### Concentration-response experiments: degraders

To evaluate the rescue effect that different IKZF1-degraders have on dCK*-IKZF1 cells under suicide substrate-induced growth inhibition, 10-point half-log serial dilutions of the degraders starting from 10 mM were prepared manually in DMSO and 20 nL of these dilutions were dispensed in quadruplicate during step 1 of the general protocol described above. In step 2, AraC was added to all wells at a final concentration of 0.3 µM (40 nL from 0.3 mM DMSO stock) and the general protocol was followed (final DMSO 0.15%).

#### Screening

In step 1 according to the general protocol, library compounds were dispensed into single wells at 5 µM final concentration (10 nL from 10 mM DMSO stock) in plate columns 3-22. Columns 2 and 24 received 10 µM pomalidomide (10 nL from 20 mM DMSO stock), and columns 1 and 23 received DMSO only (10 nL). In step 2, columns 3-24 received AraC (40 nL of 0.3 mM) and columns 1 and 2 received DMSO only (40 nL).

### Statistical analysis

#### Primary growth recovery screening

From Hoechst-based object count for each well growth (%) was determined by normalization on a plate basis to pomalidomide-treated wells (10 µM, n = 16, 100% growth) and vehicle-treated wells (DMSO, n = 16, 0% Growth). Each compound was tested in duplicate at 5 µM on different screening days and mean growth (%) was calculated. Outliers in positive and negative control wells were invalidated when exceeding 3x standard deviation of mean to increase robustness and reliability.

#### Counter screening for growth recovery

Growth (%) was determined from Hoechst-based object count by normalization on a plate basis to vehicle-treated wells that did not receive cytarabine (n = 16, 100% growth) and vehicle-treated wells that did receive cytarabine (DMSO, n = 16, 0% growth). Each drug was tested in singlicate at 5 µM. Outliers in positive and negative control wells were invalidated when exceeding 3x standard deviation of mean to increase robustness and reliability.

#### Hit detection in growth recovery format

Due to the differences in normalization for primary versus counter (simulated 100% rescue), we set different thresholds for hit detection: hits were defined as yielding signals above 10*SD_Low_ over low control in the primary (dCK*-IKZF1) and below 5*SD_Low_ over low control in the counter assay (dCK*), SD_Low_= standard deviation of the low control.

#### FRET-based screening

Inhibition (%) was determined from FRET-ratio by normalization on a plate basis to pomalidomide-treated wells (10 µM, n = 16, 100% inhibition) and vehicle-treated wells (DMSO, n = 16, 0% inhibition). Outliers in positive and negative control wells were invalidated when exceeding 3x standard deviation of mean to increase robustness and reliability.

Data were analyzed using IDBS Activity Base, KNIME and plotted using Spotfire, GraphPad Prism v10.

#### Concentration-response experiments

For determining half-maximal inhibitory concentration (IC_50_)-values for BVdU and AraC, Hoechst-based object count from wells treated with log- or half-log serial compound dilutions was normalized to averaged object counts from at least 8 wells that received DMSO vehicle instead of suicide substrate (100% growth).

For determining half-maximal inhibitory concentration (IC_50_)-values for hits from screening and additional CELMoDs in the signal rescue or signal inhibition format, each condition was tested in triplicate and normalization was done as described for screening.

Data were analyzed and plotted using GraphPad Prism v10, half-maximal inhibitory concentration (IC_50_) values were determined using the “[inhibitor] versus response -- Variable slope (four parameters)” analysis mode.

## Data availability

The supporting data for this article have been included as part of the Supporting Information.

## Associated content

### Supporting information

Additional experimental methods; signal rescue assay optimization; sensitizer TAS-114; full plasmid sequences (pdf).

Primary data files from screening; library composition including SMILES (non-blinded compounds) (xlsx).

## Supporting information

Supplementary Table: Signal Inhibition Screen

Supplementary Table: Signal Rescue Screens

Supplementary Table: Validation library

## Acknowledgements

We thank our partners in PROXIDRUGS for excellent scientific discussions and especially Prof. Felix Hausch and team for contributions to the validation library.

This work was supported by the German Federal Ministry of Education and Research (BMBF) Projects: 03ZU1109AA ‘PROXIDRUGS: ProxiDETECT’ and 03ZU1109JA ‘PROXIDRUGS: AntiDEG’.

## Author contributions

Conceptualization (JH, AZ, MAQ, PG, OP, AK), Data curation (JH, JR, AZ), Formal analysis (JH, AW, JHe, JR, AZ), Funding acquisition (MAQ, PG, OP, AK), Investigation (JH, AW, MK, MS, MB, JHe, MW), Methodology (JH, MK), Project administration (JH), Supervision (JH, PG, OP, AK), Visualization (JH, JR), Writing – original draft (JH), Writing – review & editing (all authors).

