## Supplementary Figures, Tables, Materials, Methods for "Optimization and evaluation of complementary degrader discovery assays for application in screening"

for

### Supplementary Figures and Tables

#### Assay principle and optimization of the growth recovery assay

The signal rescue assay adapts a concept from gene therapy to couple TPD to a positive readout, specifically restoration of cell growth.[1] This way, the assay only has a positive readout when cells are allowed to (partially) regain their unperturbed growth kinetics; hence, in contrast to signal inhibition-type formats, this assay inherently excludes compounds with generalized inhibitory effects such as translation inhibitors.

The assay principle relies on ectopic expression of a suicide kinase, dCK*, a triple mutant form of deoxycytidine kinase dCK, where serine 74 is replaced by glutamic acid which mimics phosphoserine, hence yielding a constitutively active kinase.[2] Moreover, arginine at position 104 is mutated to methionine and aspartic acid in position 133 is mutated to alanine, broadening the substrate spectrum towards thymidine and analogues.[3] In the gene therapy context, dCK* in conjunction with the 5-bromovinyl uridine analogue BVdU (Brivudine) is used for selective inhibition of genetically modified cells: since BVdU requires activation but does not match chemical structure requirements of cellular kinases, it is only effective in cells ectopically expressing dCK*.[4] Once BVdU is phosphorylated by dCK*, BVdU 5'-monophosphate inhibits cellular thymidylate synthase, thereby selectively stopping the growth of dCK*-positive cells (Figure S1A).

This principle was extended by Koduri *et al.* to a signal rescue-type degrader assay: by fusing dCK* to the target protein, cell proliferation can be restored via co-degradation of dCK* by a target degrader.[1]

---

[1] V. Koduri, L. Duplaquet, B.L. Lampson, et al. *Sci Adv.*, 2021, **7**:eabd6263. https://doi.org/10.1126/sciadv.abd6263
[2] S. Hazra, A. Szewczak, S. Ort, M. Konrad and A. Lavie, *Biochemistry*, 2011, **50**, 2870-2880. https://doi.org/10.1021/bi2001032
[3] S. Hazra, E. Sabini, S. Ort, M. Konrad and A. Lavie, *Biochemistry*, 2009, **48**, 1256-1263. https://doi.org/10.1038/nsb942
[4] A. Neschadim, J.C. Wang, T. Sato, D.H. Fowler, A. Lavie and J.A. Medin, *Mol. Ther.*, 2012, **20**, 1002-1013. https://doi.org/10.1038/mt.2011.298

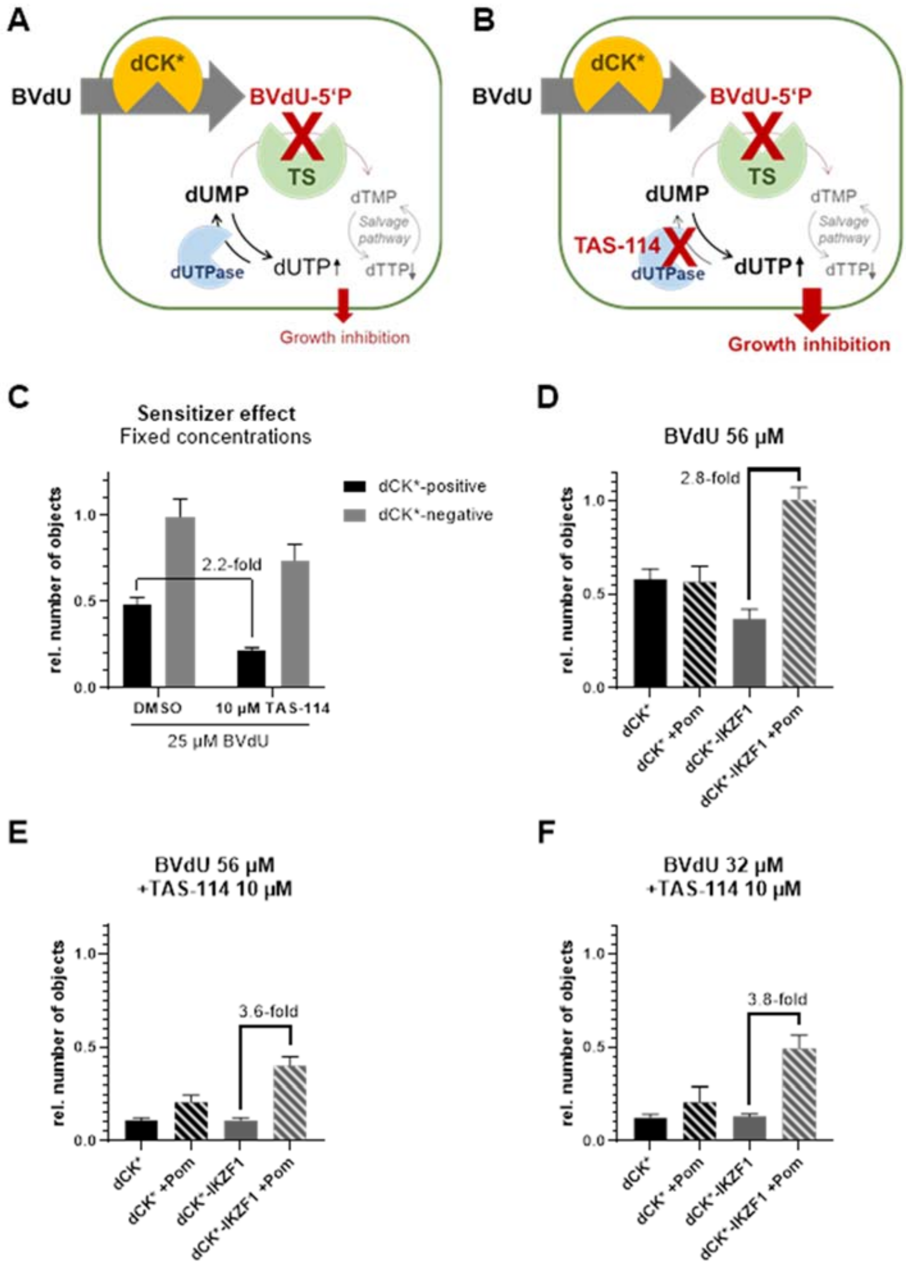

**Figure S1: BVdU mode of action and sensitizer effect of TAS-114 in BVdU-treated cells. (A)** Schematic representation of intracellular BVdU activation by dCK*. BVdU-5'P induces inhibition of cell growth via increased dUTP and concomitantly decreased dTTP level. This effect is countered by the cellular enzyme dUTPase. (**B)** Schematic representation of BVdU potency enhancement via dUTPase inhibition. **(C)** Comparison of single treatment (25 µM BVdU) and combination treatment (10 µM TAS-114 plus 25 µM BVdU) in dCK*-positive or -negative cells. p<0.000001 for 25 µM BVdU +DMSO versus 10 µM TAS-114 plus 25 µM BVdU in dCK*-positive cells (unpaired t-test). Objects represent GFP-positive cells and are retrieved via image-based counting from whole wells in 384-well plate format. Data shown are means +/- SD of 16 wells. **(D)** Growth recovery effect of 10 µM pomalidomide on dCK*- or dCK*-IKZF1-expressing cells treated with 56 µM BVdU. **(E)** Growth recovery effect of 10 µM pomalidomide on dCK*- or dCK*-IKZF1-expressing cells treated with 56 µM BVdU and 10 µM TAS-114. **(F)** Growth recovery effect of 10 µM pomalidomide on dCK*- or dCK*-IKZF1-expressing cells treated with 32 µM BVdU and 10 µM TAS-114. Number of objects are retrieved based on Hoechst staining and normalized to DMSO control without suicide substrate.

Looking to increase the assay window, we have analyzed the effect of a molecular sensitizer specific to the mode of action of BVdU. Similar to the anticancer antimetabolite 5-fluorouracil (5-FU), BVdU perturbs the cellular dNTP pool via inhibition of thymidylate synthase (TS) when metabolized intracellularly to the nucleoside analogue 5'-phosphate (BVdU-5'P).[5] More specifically, dTTP level is decreased while dUTP level is increased, ultimately leading to inhibition of cell growth. This effect is countered by cellular dUTPase (Figure S1A); hence, dUTPase inhibition (re-) sensitizes cells towards TS inhibition (Figure S1B).

We titrated TAS-114, a potent small molecule, dual dUTPase/dihydropyrimidine dehydrogenase inhibitor[6], and BVdU in dCK*-positive versus -negative cells in a matrix-type combination treatment (data not shown). As expected, increasing TAS-114 concentrations led to an enhancement of the BVdU-effect, while TAS-114 itself had no effect in this model (highest tested concentration 100 µM). Combination treatment using fixed concentrations (10 µM TAS-114 and 25 µM BVdU) resulted in a 2.2-fold (dCK*-positive) or 1.3-fold (dCK*-negative) stronger signal inhibition compared to the mono-treated (TAS-114-free) condition (Figure S1C). To test whether this sensitizer effect could increase the assay window, dCK*-IKZF1 expressing cells were pre-treated with pomalidomide or vehicle before addition of suicide substrate and TAS-114. As expected, the relative number of objects in the vehicle pre-treated wells was lower when TAS-114 was present (Figure S1D versus S1E,F). Still, the rescue effect stayed below 4-fold growth recovery by pomalidomide in BVdU-co-treated wells (Figure S1E,F).

Finally, we have optimized the signal rescue assay, especially with regard to sensitivity and robustness, by 1) switching the assessment of cell growth from GFP-based to Hoechst nuclear staining-based object counting which more faithfully represented the cell number present in the well and 2) substituting the nucleoside analogue substrate BVdU with the more effective cytarabine (AraC). AraC-5'-triphosphate serves as a substrate for cellular DNA polymerases, where it competes with canonical deoxycytidine 5'-triphosphate, and prevents further DNA synthesis and hence cell proliferation. Accordingly, its mode of action also relies on activation via phosphorylation. While, in contrast to BVdU, AraC phosphorylation is readily catalyzed by cellular deoxycytidine kinase (dCK) as well as by the triple mutant dCK*,[7] selectivity in growth inhibition was expected to be mediated by the high expression level of dCK* plus its constitutive activity versus unmodified dCK.

---

[5] J. Balzarini, E. De Clercq, A. Verbruggen, D. Ayusawa, K. Shimizu and T. Seno, *Mol Pharmacol.* 1987, **32**, 410-416. https://doi.org/10.1016/S0026-895X(25)13017-4

[6] W. Yano, T. Yokogawa, T. Wakasa, et al. *Mol Cancer Ther.* 2018, **17**, 1683-1693. https://doi.org/10.1158/1535-7163.mct-17-0911

[7] S. Hazra, S. Ort, M. Konrad and A. Lavie, *Biochemistry*, 2010, **49**, 6784-6790. https://doi.org/10.1021/bi100839e

#### TPD validation library and statistical analysis of hits from the library subsets

This library was designed with the aim to assess the individual strengths and limitations of the different TPD screening assays. It leverages prior knowledge of the compounds' biological effects as well as (predicted) compound promiscuity,[8] i.e., frequent activity in a range of biological contexts, thus pointing towards target-unrelated effects. Such promiscuity may result from poly-pharmacology of a compound or pharmacological activity at central nodes in cellular pathways leading to more generalized downstream effects (e.g., translation inhibitors). It includes a subset of blinded proprietary compounds with well-understood bioactivities that should in general not fall within the class of predicted frequent hitters. Class 1 and 2 compound SMILES are included in supplementary xlsx-file.

**Supplementary Table 1. Composition of TPD validation library**

| | Class | # of compounds | proportion |
|---|---|---|---|
| **Total compounds** | 1-3 | 941 | |
| **Known bioactives** (incl. degraders) | 1 | 305 | 0.32 |
| **Frequent hitters** | 2 | 324 | 0.34 |
| **Blinded compounds** | 3 | 312 | 0.33 |

**Supplementary Table 2. Comparative statistical analysis of hits from the validation library** by subset in signal inhibition and signal rescue assay format.

| | | **Growth recovery** (primary) | | | **Growth recovery** (primary + counter) | | | **Growth inhibitor** | | | **FRET signal inhibition** | | |
|---|---|---|---|---|---|---|---|---|---|---|---|---|---|
| | | # | proportion | rate | # | proportion | rate | # | proportion | rate | # | proportion | rate |
| **total hits** | | 17 | | 1.81% | 7 | | 0.74% | 156 | | 16.58% | 45 | | 4.78% |
| **blinded hits** | | 4 | 0.24 | 0.43% | 3 | 0.43 | 0.32% | 28 | 0.18 | 2.98% | 9 | 0.20 | 0.96% |
| **non-blinded hits** | FH | 1 | 0.06 | 0.11% | 0 | 0.00 | 0.00% | 102 | 0.65 | 10.84% | 23 | 0.51 | 2.44% |
| | non-FH | 12 | 0.71 | 1.28% | 4 | 0.57 | 0.43% | 26 | 0.17 | 2.76% | 13 | 0.29 | 1.38% |

### is number of compounds; FH is frequent hitter as predicted via the HitDexter machine learning model;[8] rate is defined as 100*(#/Total number of compounds in library); compounds are considered as inhibitors in the growth recovery format when resulting in <-11% growth.

[8] C. Stork, Y. Chen, M. Šícho and J. Kirchmair, *J. Chem. Inf. Model.*, 2019, **59**, 1030-1043. nerdd.univie.ac.at/hitdexter/

#### Supplementary Materials and Methods

##### Commercial material used in this study

| **Material** | **Supplier** | **Cat.-No.** |
|---|---|---|
| Cell culture | | |
| HEK293FT cells | Life Technologies GmbH | R70007 |
| Phosphate buffered saline (PBS; Gibco DPBS w/o calcium, magnesium) | Life Technologies GmbH | 14190250 |
| DMEM (high glucose, no L-glutamine, no phenol red) | Life Technologies GmbH | 31053044 |
| Fetal bovine serum (FBS) | Capricorn GmbH | 10-FBS-11F |
| L-Glutamine (Gibco) | Life Technologies GmbH | 25030081 |
| Penicillin/streptomycin (100 U/mL penicillin, and 100 mg/mL streptomycin, Gibco) | Life Technologies GmbH | 11140035 |
| MEM non-essential amino acids (Gibco) | Life Technologies GmbH | 11140035 |
| Sodium pyruvate (Gibco) | Life Technologies GmbH | 11360070 |
| Trypsin-EDTA 0.05% | Capricorn GmbH | TRY-1B |
| G-418 sulfate, 50 mg/mL | Capricorn GmbH | G418-B |
| Blasticidin | Life Technologies GmbH | R21001 |
| Compounds | | |
| 5-Bromovinyluridine (BVdU) | TCI Deutschland GmbH | B3404 |
| Cytarabine (AraC) | Sigma-Aldrich Chemie GmbH | C6645 |
| Hoechst 33342 | Merck Chemicals GmbH | B2261 |
| TAS-114 | MedChemExpress | HY-124062 |
| Pomalidomide | MedChemExpress | HY-10984 |
| Iberdomide | MedChemExpress | HY-101291 |
| Avadomide | MedChemExpress | HY-100507 |
| Mezigdomide | MedChemExpress | HY-129395 |
| NVP-DKY709 | MedChemExpress | HY-144998 |
| Eragidomide | MedChemExpress | HY-130800 |
| MRT-2359 | MedChemExpress | HY-153356 |
| CC-885 | MedChemExpress | HY-101488 |
| Thalidomide 5-fluoride | MedChemExpress | HY-W087383 |

#### Generation of modified HEK293FT cell lines for the signal-rescue assay

*Production of lentiviral particles*

For the production of lentiviral particles, the protocol described by Koduri *et al.* was followed.[1] Briefly, HEK293FT cells were detached using Accutase and 20x10$^6$ cells were seeded in a 10 cm dish in 10 mL complete medium (without selection antibiotics) for cells to reach 90-95% confluency the next day. Then, medium was replaced with 5 mL fresh complete medium and lipofectamine 2000-based cotransfection was performed with packaging (psPAX2), envelope (pMD2.G) and transfer (pLX304-based) plasmid in a ratio of 5:1:5 in a total of 3 mL Opti-MEM medium (serum-free; total amounts used were 7.7 µg DNA and 25 µL Lipofectamine). The next day, medium was replaced by 5 mL fresh complete medium and after another 24 h, media was harvested and stored at 4 °C, and 5 mL fresh complete medium were added to the lentiviral particle-producing cells. Media was again harvested at 72 h post transfection, combined with the first harvest and centrifuged at 3000 rpm for 15 min at 4 °C. Supernatant was filtered through 0.45 µm PES filter and directly used for transduction.

*Transduction and selection of cells*

For the generation of stably modified cells, the protocol by Koduri *et al.* was followed.[1] Briefly, HEK293FT cells were detached using Accutase and 6x10$^6$ cells were seeded in a 10 cm dish in 5 mL complete medium (without selection antibiotics). After allowing cells to settle for 5 h, 5 mL of medium containing lentiviral particles (1 mL of harvest medium + 4 mL fresh complete medium) were supplemented with 8 µg/mL polybrene and added to the cells. The next day, medium was replaced by fresh complete medium. Cells were selected by growth in complete medium supplemented with 10 µg/mL Blasticidin and 800 µg/mL G-418 starting at 48 h after transduction. Cells were grown in T75 cell culture flasks, medium was changed every 1-2 days and cells were passed based on need to keep confluency below 80% (~every 3-4 days) for 2 weeks. To assess for any residual lentiviral particles, qPCR (HIV-1-pol-1 and VSV-G) on cell culture supernatant was shown negative after a total of 10 medium changes and 3 passages. After a total of 10 passages, cultures were grown in complete medium supplemented with 8 µg/mL Blasticidin and 800 µg/mL G-418. Expression of tagged protein of correct size was confirmed by immunoblot analysis (V5). Flow cytometry for dCK*-only and dCK*-IKZF1 cells (GFP) confirmed >95% GFP-positive population. IKZF1-modified cells were only at 77% GFP-positive and were hence sorted for 1% highest expressers.

#### Plasmids

*Generation of lentiviral expression vectors (transfer plasmids)*

Lentiviral expression vectors were a kind gift from the Kaelin lab at Harvard;[1] since sequencing revealed some instabilities, plasmids were re-cloned using the uninterrupted coding sequences from these original vectors as templates.

After cloning, all plasmids were validated by overlapping sequencing.

All plasmids contained the following markers for resistance:

Bacterial culture: ampicillin

Cell culture selection marker: blasticidin

##### pLX304-IKZF1-IRES-GFP

Bicistronic lentiviral vector with IRES (internal ribosomal binding site) enabling CMV controlled expression of IKZF1 protein (isoform Ik7; UniProt ID Q13422-7), and eGFP (enhanced GFP).

Expressed protein: IKZF1(Ik7)-V5

```
1        10        20        30        40        50
|        |         |         |         |         |
MDADEGQDMSQVSGKESPPVSDTPDEGDEPMPIPEDLSTTSGGQQSSKSD
RVVASNVKVETQSDEENGRACEMNGEECAEDLRMLDASGEKMNGSHRDQG
SSALSGVGGIRLPNGKLKCDICGIICIGPNVLMVHKRSHTGERPFQCNQC
GASFTQKGNLLRHIKLHSGEKPFKCHLCNYACRRRDALTGHLRTHSVIKE
ETNHSEMAEDLCKIGSERSLVLDRLASNVAKRKSSMPQKFLGDKGLSDTP
YDSSASYEKENEMMKSHVMDQAINNAINYLGAESLRPLVQTPPGGSEVVP
VISPMYQLHKPLAEGTPRSNHSAQDSAVENLLLLSKAKLVPSEREASPSN
SCQDSTDTESNNEEQRSGLIYLTNHIAPHARNGLSLKEEHRAYDLLRAAS
ENSQDALRVVSTSGEQMKVYKCEHCRVLFLDHVMYTIHMGCHGFRDPFEC
NMCGYHSQDRYEFSSHITRGEHRFHMSNPAFLYKVVGKPIPNPLLGLDST
```

Expressed fluorescent marker: eGFP (enhanced GFP):

```
1        10        20        30        40        50
|        |         |         |         |         |
MVSKGEELFTGVVPILVELDGDVNGHKFSVSGEGEGDATYGKLTLKFICT
TGKLPVPWPTLVTTLTYGVQCFSRYPDHMKQHDFFKSAMPEGYVQERTIF
FKDDGNYKTRAEVKFEGDTLVNRIELKGIDFKEDGNILGHKLEYNYNSHN
VYIMADKQKNGIKVNFKIRHNIEDGSVQLADHYQQNTPIGDGPVLLPDNH
YLSTQSALSKDPNEKRDHMVLLEFVTAAGITLGMDELYK
```

Full plasmid sequence:

```
  1 gcccggggtt attaatagta atcaattacg gggtcattag ttcatagccc atatatggag
 61 ttccgcgtta cataacttac ggtaaatggc ccgcctggct gaccgcccaa cgacccccgc
121 ccattgacgt caataatgac gtatgttccc atagtaacgc caatagggac tttccattga
181 cgtcaatggg tggagtattt acggtaaact gcccacttgg cagtacatca agtgtatcat
241 atgccaagta cgccccctat tgacgtcaat gacggtaaat ggcccgcctg gcattatgcc
301 cagtacatga ccttatggga ctttcctact tggcagtaca tctacgtatt agtcatcgct
361 attaccatgg tgatgcggtt ttggcagtac atcaatgggc gtggatagcg gtttgactca
421 cggggatttc caagtctcca ccccattgac gtcaatggga gtttgttttg gcaccaaaat
481 caacggactt tccaaaatgt cgtaacaact ccgccccatt gacgcaaatg ggcggtaggc
```

541 cgtgtacggt gggaggtcta tataagcaga gctctctggc taactgtcgg gatcaacaag
601 tttgtacaaa aaagttggca tggatgctga tgagggtcaa gacatgtccc aagtttcagg
661 gaaggaaagc cccctgtaa gcgatactcc agatgagggc gatgagccca tgccgatccc
721 cgaggacctc tccaccacct cgggaggaca gcaaagctcc aagagtgaca gagtcgtggc
781 cagtaatgtt aaagtagaga ctcagagtga tgaagagaat gggcgtgcct gtgaaatgaa
841 tggggaagaa tgtgcggagg atttacgaat gcttgatgcc tcgggagaga aaatgaatgg
901 ctcccacagg gaccaaggca gctcggcttt gtcgggagtt ggaggcattc gacttcctaa
961 cggaaaacta aagtgtgata tctgtgggat catttgcatc gggcccaatg tgctcatggt
1021 tcacaaaaga agccacactg gagaacggcc cttccagtgc aatcagtgcg gggcctcatt
1081 cacccagaag ggcaacctgc tccggcacat caagctgcat tccggggaga agcccttcaa
1141 atgccacctc tgcaactacg cctgccgccg gagggacgcc ctcactggcc acctgaggac
1201 gcactccgtc attaaagaag aaactaatca cagtgaaatg gcagaagacc tgtgcaagat
1261 aggatcagag agatctctcg tgctggacag actagcaagt aacgtcgcca aacgtaagag
1321 ctctatgcct cagaaatttc ttggggacaa gggcctgtcc gacacgccct acgacagcag
1381 cgccagctac gagaaggaga acgaaatgat gaagtcccac gtgatggacc aagccatcaa
1441 caacgccatc aactacctgg gggccgagtc cctgcgcccg ctggtgcaga cgcccccggg
1501 cggttccgag gtggtcccgg tcatcagccc gatgtaccag ctgcacaagc cgctcgcgga
1561 gggcaccccg cgctccaacc actcggccca ggacagcgcc gtggagaacc tgctgctgct
1621 ctccaaggcc aagttggtgc cctcggagcg cgaggcgtcc ccgagcaaca gctgccaaga
1681 ctccacggac accgagagca acaacgagga gcagcgcagc ggtctcatct acctgaccaa
1741 ccacatcgcc ccgcacgcgc gcaacgggct gtcgctcaag gaggagcacc gcgcctacga
1801 cctgctgcgc gccgcctccg agaactcgca ggacgcgctc cgcgtggtca gcaccagcgg
1861 ggagcagatg aaggtgtaca agtgcgaaca ctgccgggtg ctcttcctgg atcacgtcat
1921 gtacaccatc cacatgggct gccacggctt ccgtgatcct tttgagtgca acatgtgcgg
1981 ctaccacagc caggaccggt acgagttctc gtcgcacata acgcgagggg agcaccgctt
2041 ccacatgagc aacccagctt tcttgtacaa agtggttggt aagcctatcc ctaaccctct
2101 cctcggtctc gattctacgt agtaatgagc tagccgctac gtaaattccg cccccccccc
2161 ccctctccct cccccccccc taacgttact ggccgaagcc gcttggaata aggccggtgt
2221 gcgtttgtct atatgttatt ttccaccata ttgccgtctt ttggcaatgt gagggcccgg
2281 aaacctggcc ctgtcttctt gacgagcatt cctaggggtc tttcccctct cgccaaagga
2341 atgcaaggtc tgttgaatgt cgtgaaggaa gcagttcctc tggaagcttc ttgaagacaa
2401 acaacgtctg tagcgaccct ttgcaggcag cggaaccccc cacctggcga caggtgcctc
2461 tgcggccaaa agccacgtgt ataagataca cctgcaaagg cggcacaacc ccagtgccac
2521 gttgtgagtt ggatagttgt ggaaagagtc aaatggctct cctcaagcgt attcaacaag
2581 gggctgaagg atgcccagaa ggtaccccat tgtatgggat ctgatctggg gcctcggtgc
2641 acatgcttta catgtgttta gtcgaggtta aaaaaacgtc taggcccccc gaaccacggg
2701 gacgtggttt tcctttgaaa aacacgatga taatatggcc acaaccatgg tgagcaaggg
2761 cgaggagctg ttcaccgggg tggtgcccat cctggtcgag ctggacggcg acgtaaacgg
2821 ccacaagttc agcgtgtccg gcgagggcga gggcgatgcc acctacggca agctgaccct
2881 gaagttcatc tgcaccaccg gcaagctgcc cgtgccctgg cccaccctcg tgaccaccct
2941 gacctacggc gtgcagtgct tcagccgcta ccccgaccac atgaagcagc acgacttctt
3001 caagtccgcc atgcccgaag gctacgtcca ggagcgcacc atcttcttca aggacgacgg
3061 caactacaag acccgcgccg aggtgaagtt cgagggcgac accctggtga accgcatcga
3121 gctgaagggc atcgacttca aggaggacgg caacatcctg gggcacaagc tggagtacaa
3181 ctacaacagc cacaacgtct atatcatggc cgacaagcag aagaacggca tcaaggtgaa
3241 cttcaagatc cgccacaaca tcgaggacgg cagcgtgcag ctcgccgacc actaccagca
3301 gaacaccccc atcggcgacg gccccgtgct gctgcccgac aaccactacc tgagcaccca
3361 gtccgccctg agcaaagacc ccaacgagaa gcgcgatcac atggtcctgc tggagttcgt
3421 gaccgccgcc gggatcactc tcggcatgga cgagctgtac aagtaaaccg gtggcgcgtt
3481 aagtcgacaa tcaacctctg gattacaaaa tttgtgaaag attgactggt attcttaact
3541 atgttgctcc ttttacgcta tgtggatacg ctgctttaat gcctttgtat catgctattg
3601 cttcccgtat ggctttcatt ttctcctcct tgtataaatc ctggttgctg tctctttatg
3661 aggagttgtg gcccgttgtc aggcaacgtg gcgtggtgtg cactgtgttt gctgacgcaa
3721 cccccactgg ttggggcatt gccaccacct gtcagctcct ttccgggact ttcgctttcc
3781 ccctccctat tgccacggcg gaactcatcg ccgcctgcct tgcccgctgc tggacagggg
3841 ctcggctgtt gggcactgac aattccgtgg tgttgtcggg gaaatcatcg tcctttcctt
3901 ggctgctcgc ctgtgttgcc acctggattc tgcgcgggac gtccttctgc tacgtccctt
3961 cggccctcaa tccagcggac cttccttccc gcggcctgct gccggctctg cggcctcttc
4021 cgcgtcttcg ccttcgccct cagacgagtc ggatctccct ttgggccgcc tccccgcgtc

4081 gactttaaga ccaatgactt acaaggcagc tgtagatctt agccactttt taaaagaaaa
4141 ggggggactg gaagggctaa ttcactccca acgaagacaa gatctgcttt ttgcttgtac
4201 tgggtctctc tggttagacc agatctgagc ctgggagctc tctggctaac tagggaaccc
4261 actgcttaag cctcaataaa gcttgccttg agtgcttcaa gtagtgtgtg cccgtctgtt
4321 gtgtgactct ggtaactaga gatccctcag acccttttag tcagtgtgga aaatctctag
4381 cagtacgtat agtagttcat gtcatcttat tattcagtat ttataacttg caaagaaatg
4441 aatatcagag agtgagagga acttgtttat tgcagcttat aatggttaca aataaagcaa
4501 tagcatcaca aatttcacaa ataaagcatt tttttcactg cattctagtt gtggtttgtc
4561 caaactcatc aatgtatctt atcatgtctg gctctagcta tcccgcccct aactccgccc
4621 atcccgcccc taactccgcc cagttccgcc cattctccgc cccatggctg actaattttt
4681 tttatttatg cagaggccga ggccgcctcg gcctctgagc tattccagaa gtagtgagga
4741 ggcttttttg gaggcctagg gacgtaccca attcgcccta tagtgagtcg tattacgcgc
4801 gctcactggc cgtcgtttta caacgtcgtg actgggaaaa ccctggcgtt acccaactta
4861 atcgccttgc agcacatccc cctttcgcca gctggcgtaa tagcgaagag gcccgcaccg
4921 atcgcccttc ccaacagttg cgcagcctga atggcgaatg ggacgcgccc tgtagcggcg
4981 cattaagcgc ggcgggtgtg gtggttacgc gcagcgtgac cgctacactt gccagcgccc
5041 tagcgcccgc tcctttcgct ttcttccctt cctttctcgc cacgttcgcc ggctttcccc
5101 gtcaagctct aaatcggggg ctccctttag ggttccgatt tagtgcttta cggcacctcg
5161 accccaaaaa acttgattag ggtgatggtt cacgtagtgg gccatcgccc tgatagacgg
5221 tttttcgccc tttgacgttg gagtccacgt tctttaatag tggactcttg ttccaaactg
5281 gaacaacact caaccctatc tcggtctatt cttttgattt ataagggatt ttgccgattt
5341 cggcctattg gttaaaaaat gagctgattt aacaaaaatt taacgcgaat tttaacaaaa
5401 tattaacgct tacaatttag gtggcacttt tcggggaaat gtgcgcggaa cccctatttg
5461 tttatttttc taaatacatt caaatatgta tccgctcatg agacaataac cctgataaat
5521 gcttcaataa tattgaaaaa ggaagagtat gagtattcaa catttccgtg tcgcccttat
5581 tccctttttt gcggcatttt gccttcctgt ttttgctcac ccagaaacgc tggtgaaagt
5641 aaaagatgct gaagatcagt tgggtgcacg agtgggttac atcgaactgg atctcaacag
5701 cggtaagatc cttgagagtt ttcgccccga agaacgtttt ccaatgatga gcacttttaa
5761 agttctgcta tgtggcgcgg tattatcccg tattgacgcc gggcaagagc aactcggtcg
5821 ccgcatacac tattctcaga atgacttggt tgagtactca ccagtcacag aaaagcatct
5881 tacggatggc atgacagtaa gagaattatg cagtgctgcc ataaccatga gtgataacac
5941 tgcggccaac ttacttctga caacgatcgg aggaccgaag gagctaaccg cttttttgca
6001 caacatgggg gatcatgtaa ctcgccttga tcgttgggaa ccggagctga atgaagccat
6061 accaaacgac gagcgtgaca ccacgatgcc tgtagcaatg gcaacaacgt tgcgcaaact
6121 attaactggc gaactactta ctctagcttc ccggcaacaa ttaatagact ggatggaggc
6181 ggataaagtt gcaggaccac ttctgcgctc ggcccttccg gctggctggt ttattgctga
6241 taaatctgga gccggtgagc gtgggtctcg cggtatcatt gcagcactgg ggccagatgg
6301 taagccctcc cgtatcgtag ttatctacac gacggggagt caggcaacta tggatgaacg
6361 aaatagacag atcgctgaga taggtgcctc actgattaag cattggtaac tgtcagacca
6421 agtttactca tatatacttt agattgattt aaaacttcat ttttaattta aaaggatcta
6481 ggtgaagatc ctttttgata atctcatgac caaaatccct taacgtgagt tttcgttcca
6541 ctgagcgtca gaccccgtag aaaagatcaa aggatcttct tgagatcctt tttttctgcg
6601 cgtaatctgc tgcttgcaaa caaaaaaacc accgctacca gcggtggttt gtttgccgga
6661 tcaagagcta ccaactcttt ttccgaaggt aactggcttc agcagagcgc agataccaaa
6721 tactgttctt ctagtgtagc cgtagttagg ccaccacttc aagaactctg tagcaccgcc
6781 tacatacctc gctctgctaa tcctgttacc agtggctgct gccagtggcg ataagtcgtg
6841 tcttaccggg ttggactcaa gacgatagtt accggataag gcgcagcggt cgggctgaac
6901 ggggggttcg tgcacacagc ccagcttgga gcgaacgacc tacaccgaac tgagatacct
6961 acagcgtgag ctatgagaaa gcgccacgct tcccgaaggg agaaaggcgg acaggtatcc
7021 ggtaagcggc agggtcggaa caggagagcg cacgagggag cttccagggg gaaacgcctg
7081 gtatctttat agtcctgtcg ggtttcgcca cctctgactt gagcgtcgat ttttgtgatg
7141 ctcgtcaggg gggcggagcc tatggaaaaa cgccagcaac gcggcctttt tacggttcct
7201 ggccttttgc tggccttttg ctcacatgtt ctttcctgcg ttatcccctg attctgtgga
7261 taaccgtatt accgcctttg agtgagctga taccgctcgc cgcagccgaa cgaccgagcg
7321 cagcgagtca gtgagcgagg aagcggaaga gcgcccaata cgcaaaccgc ctctccccgc
7381 gcgttggccg attcattaat gcagctggca cgacaggttt cccgactgga aagcgggcag
7441 tgagcgcaac gcaattaatg tgagttagct cactcattag gcaccccagg ctttacactt
7501 tatgcttccg gctcgtatgt tgtgtggaat tgtgagcgga taacaatttc acacaggaaa
7561 cagctatgac catgattacg ccaagcgcgc aattaaccct cactaaaggg aacaaaagct

```
 7621 ggagctgcaa gcttaatgta gtcttatgca atactcttgt agtcttgcaa catggtaacg
 7681 atgagttagc aacatgcctt acaaggagag aaaaagcacc gtgcatgccg attggtggaa
 7741 gtaaggtggt acgatcgtgc cttattagga aggcaacaga cgggtctgac atggattgga
 7801 cgaaccactg aattgccgca ttgcagagat attgtattta agtgcctagc tcgatacata
 7861 aacgggtctc tctggttaga ccagatctga gcctgggagc tctctggcta actagggaac
 7921 ccactgctta agcctcaata aagcttgcct tgagtgcttc aagtagtgtg tgcccgtctg
 7981 ttgtgtgact ctggtaacta gagatccctc agaccctttt agtcagtgtg gaaaatctct
 8041 agcagtggcg cccgaacagg gacttgaaag cgaaagggaa accagaggag ctctctcgac
 8101 gcaggactcg gcttgctgaa gcgcgcacgg caagaggcga ggggcggcga ctggtgagta
 8161 cgccaaaaat tttgactagc ggaggctaga aggagagaga tgggtgcgag agcgtcagta
 8221 ttaagcgggg gagaattaga tcgcgatggg aaaaaattcg gttaaggcca gggggaaaga
 8281 aaaaatataa attaaaacat atagtatggg caagcaggga gctagaacga ttcgcagtta
 8341 atcctggcct gttagaaaca tcagaaggct gtagacaaat actgggacag ctacaaccat
 8401 cccttcagac aggatcagaa gaacttagat cattatataa tacagtagca accctctatt
 8461 gtgtgcatca aaggatagag ataaaagaca ccaaggaagc tttagacaag atagaggaag
 8521 agcaaaacaa aagtaagacc accgcacagc aagcggccgc tgatcttcag acctggagga
 8581 ggagatatga gggacaattg gagaagtgaa ttatataaat ataaagtagt aaaaattgaa
 8641 ccattaggag tagcacccac caaggcaaag agaagagtgg tgcagagaga aaaaagagca
 8701 gtgggaatag gagctttgtt ccttgggttc ttgggagcag caggaagcac tatgggcgca
 8761 gcgtcaatga cgctgacggt acaggccaga caattattgt ctggtatagt gcagcagcag
 8821 aacaatttgc tgagggctat tgaggcgcaa cagcatctgt tgcaactcac agtctggggc
 8881 atcaagcagc tccaggcaag aatcctggct gtggaaagat acctaaagga tcaacagctc
 8941 ctggggattt ggggttgctc tggaaaactc atttgcacca ctgctgtgcc ttggaatgct
 9001 agttggagta ataaatctct ggaacagatt tggaatcaca cgacctggat ggagtgggac
 9061 agagaaatta acaattacac aagcttaata cactccttaa ttgaagaatc gcaaaaccag
 9121 caagaaaaga atgaacaaga attattggaa ttagataaat gggcaagttt gtggaattgg
 9181 tttaacataa caaattggct gtggtatata aaattattca taatgatagt aggaggcttg
 9241 gtaggtttaa gaatagtttt tgctgtactt tctatagtga atagagttag gcagggatat
 9301 tcaccattat cgtttcagac ccacctccca accccgaggg gacccttgcg cctttttccaa
 9361 ggcagccctg ggtttgcgca gggacgcggc tgctctgggc gtggttccgg gaaacgcagc
 9421 ggcgccgacc ctgggtctcg cacattcttc acgtccgttc gcagcgtcac ccggatcttc
 9481 gccgctaccc ttgtgggccc cccggcgacg cttcctgctc cgcccctaag tcgggaaggt
 9541 tccttgcggt tcgcggcgtg ccggacgtga caaacggaag ccgcacgtct cactagtacc
 9601 ctcgcagacg gacagcgcca gggagcaatg gcagcgcgcc gaccgcgatg ggctgtggcc
 9661 aatagcggct gctcagcagg gcgcgccgag agcagcggcc gggaaggggc ggtgcgggag
 9721 gcggggtgtg gggcggtagt gtgggccctg ttcctgcccg cgcggtgttc cgcattctgc
 9781 aagcctccgg agcgcacgtc ggcagtcggc tccctcgttg accgaatcac cgacctctct
 9841 ccccaggggg taccaccatg gccaagcctt tgtctcaaga agaatccacc ctcattgaaa
 9901 gagcaacggc tacaatcaac agcatcccca tctctgaaga ctacagcgtc gccagcgcag
 9961 ctctctctag cgacggccgc atcttcactg gtgtcaatgt atatcatttt actgggggac
10021 cttgtgcaga actcgtggtg ctgggcactg ctgctgctgc ggcagctggc aacctgactt
10081 gtatcgtcgc gatcggaaat gagaacaggg gcatcttgag cccctgcgga cggtgccgac
10141 aggtgcttct cgatctgcat cctgggatca aagccatagt gaaggacagt gatggacagc
10201 cgacggcagt tgggattcgt gaattgctgc cctctggtta tgtgtgggag ggcctgcagc
10261 tgcagtagta agaattctag atcttgagac aaatggcagt attcatccac aattttaaaa
10321 gaaaaggggg gattgggggg tacagtgcag gggaaagaat agtagacata atagcaacag
10381 acatacaaac taaagaatta caaaaacaaa ttacaaaaat tcaaaatttt cgggtttatt
10441 acagggacag cagagatcca ctttggcgcc ggctcgaggg g
//
```

pLX304-DCK*-IRES-GFP

Bicistronic lentiviral vector with IRES (internal ribosomal binding site) enabling CMV controlled expression of DCK* protein, and eGFP (enhanced GFP).

DCK* encompasses three mutations in comparison to native *h.s.* deoxycytidine kinase (UniProt ID P27707): S74E; R104M; D133A.

Expressed protein: DCK*-V5

```
1         10        20        30        40        50
|         |         |         |         |         |
MVPRGSHMATPPKRSCPSFSASSEGTRIKKISIEGNIAAGKSTFVNILKQ
LCEDWEVVPEPVARWCNVQSTQDEFEELTMEQKNGGNVLQMMYEKPERWS
FTFQTYACLSMIRAQLASLNGKLKDAEKPVLFFERSVYSARYIFASNLYE
SECMNETEWTIYQDWHDWMNNQFGQSLELDGIIYLQATPETCLHRIYLRG
RNEEQGIPLEYLEKLHYKHESWLLHRTLKTNFDYLQEVPILTLDVNEDFK
DKYESLVEKVKEFLSTLNPAFLYKVVGKPIPNPLLGLDST
```

Expressed fluorescent marker: eGFP (enhanced GFP):

```
1         10        20        30        40        50
|         |         |         |         |         |
MVSKGEELFTGVVPILVELDGDVNGHKFSVSGEGEGDATYGKLTLKFICT
TGKLPVPWPTLVTTLTYGVQCFSRYPDHMKQHDFFKSAMPEGYVQERTIF
FKDDGNYKTRAEVKFEGDTLVNRIELKGIDFKEDGNILGHKLEYNYNSHN
VYIMADKQKNGIKVNFKIRHNIEDGSVQLADHYQQNTPIGDGPVLLPDNH
YLSTQSALSKDPNEKRDHMVLLEFVTAAGITLGMDELYK
```

Full plasmid sequence:

```
   1 gcccggggtt attaatagta atcaattacg gggtcattag ttcatagccc atatatggag
  61 ttccgcgtta cataacttac ggtaaatggc ccgcctggct gaccgcccaa cgacccccgc
 121 ccattgacgt caataatgac gtatgttccc atagtaacgc caatagggac tttccattga
 181 cgtcaatggg tggagtattt acggtaaact gcccacttgg cagtacatca agtgtatcat
 241 atgccaagta cgccccctat tgacgtcaat gacggtaaat ggcccgcctg gcattatgcc
 301 cagtacatga ccttatggga ctttcctact tggcagtaca tctacgtatt agtcatcgct
 361 attaccatgg tgatgcggtt ttggcagtac atcaatgggc gtggatagcg gtttgactca
 421 cggggatttc caagtctcca ccccattgac gtcaatggga gtttgttttg gcaccaaaat
 481 caacgggact ttccaaaatg tcgtaacaac tccgccccat tgacgcaaat gggcggtagg
 541 cgtgtacggt gggaggtcta tataagcaga gctctctggc taagccacca tggttccgcg
 601 tggctctcat atggccaccc cgcccaagag aagctgccca tctttctcag ccagctctga
 661 ggggacccgc atcaagaaaa tctccatcga agggaacatc gctgcaggga agtcaacatt
 721 tgtgaatatc cttaaacaat tgtgtgaaga ttgggaagtg gttcctgaac ctgttgccag
 781 atggtgcaat gttcaaagta ctcaagatga atttgaggaa cttacaatgg agcagaaaaa
 841 tggtgggaat gttcttcaga tgatgtatga gaaacctgaa cgatggtctt ttaccttcca
 901 aacctacgcc tgtctcagta tgataagagc tcagcttgcc tctctgaatg gcaagctcaa
 961 agatgcagag aaacctgtat tatttttga acgatctgtg tatagtgcga ggtatatttt
1021 tgcatctaat ttgtatgaat ctgaatgcat gaatgagaca gagtggacaa tttatcaaga
1081 ctggcatgac tggatgaata accaatttgg ccaaagcctt gaattggatg gaatcattta
1141 tcttcaagcc actccagaga catgcttaca tagaatatat ttacggggaa gaaatgaaga
1201 gcaaggcatt cctcttgaat atttagagaa gcttcattat aaacatgaaa gctggctcct
1261 gcataggaca ctgaaaacca acttcgatta tcttcaagag gtgcctatct taacactgga
1321 tgttaatgaa gactttaaag acaaatatga aagtctggtt gaaaaggtca aagagttttt
1381 gagtactttg aacccagctt tcttgtacaa agtggttggt aagcctatcc ctaaccctct
1441 cctcggtctc gattctacgt agtaatgagc tagccgctac gtaaattccg cccccccccc
1501 ccctctccct cccccccccc taacgttact ggccgaagcc gcttggaata aggccggtgt
```

1561 gcgtttgtct atatgttatt ttccaccata ttgccgtctt ttggcaatgt gagggcccgg
1621 aaacctggcc ctgtcttctt gacgagcatt cctaggggtc tttcccctct cgccaaagga
1681 atgcaaggtc tgttgaatgt cgtgaaggaa gcagttcctc tggaagcttc ttgaagacaa
1741 acaacgtctg tagcgaccct ttgcaggcag cggaaccccc cacctggcga caggtgcctc
1801 tgcggccaaa agccacgtgt ataagataca cctgcaaagg cggcacaacc ccagtgccac
1861 gttgtgagtt ggatagttgt ggaaagagtc aaatggctct cctcaagcgt attcaacaag
1921 gggctgaagg atgcccagaa ggtaccccat tgtatgggat ctgatctggg gcctcggtgc
1981 acatgcttta catgtgttta gtcgaggtta aaaaaacgtc taggcccccc gaaccacggg
2041 gacgtggttt tcctttgaaa aacacgatga taatatggcc acaaccatgg tgagcaaggg
2101 cgaggagctg ttcaccgggg tggtgcccat cctggtcgag ctggacggcg acgtaaacgg
2161 ccacaagttc agcgtgtccg gcgagggcga gggcgatgcc acctacggca agctgaccct
2221 gaagttcatc tgcaccaccg gcaagctgcc cgtgccctgg cccaccctcg tgaccaccct
2281 gacctacggc gtgcagtgct tcagccgcta ccccgaccac atgaagcagc acgacttctt
2341 caagtccgcc atgcccgaag gctacgtcca ggagcgcacc atcttcttca aggacgacgg
2401 caactacaag acccgcgccg aggtgaagtt cgagggcgac accctggtga accgcatcga
2461 gctgaagggc atcgacttca aggaggacgg caacatcctg gggcacaagc tggagtacaa
2521 ctacaacagc cacaacgtct atatcatggc cgacaagcag aagaacggca tcaaggtgaa
2581 cttcaagatc cgccacaaca tcgaggacgg cagcgtgcag ctcgccgacc actaccagca
2641 gaacaccccc atcggcgacg gccccgtgct gctgcccgac aaccactacc tgagcaccca
2701 gtccgccctg agcaaagacc ccaacgagaa gcgcgatcac atggtcctgc tggagttcgt
2761 gaccgccgcc gggatcactc tcggcatgga cgagctgtac aagtaaaccg gtggcgcgtt
2821 aagtcgacaa tcaacctctg gattacaaaa tttgtgaaag attgactggt attcttaact
2881 atgttgctcc ttttacgcta tgtggatacg ctgctttaat gcctttgtat catgctattg
2941 cttcccgtat ggctttcatt ttctcctcct tgtataaatc ctggttgctg tctctttatg
3001 aggagttgtg gcccgttgtc aggcaacgtg gcgtggtgtg cactgtgttt gctgacgcaa
3061 cccccactgg ttggggcatt gccaccacct gtcagctcct ttccgggact ttcgctttcc
3121 ccctccctat tgccacggcg gaactcatcg ccgcctgcct tgcccgctgc tggacagggg
3181 ctcggctgtt gggcactgac aattccgtgg tgttgtcggg gaaatcatcg tcctttcctt
3241 ggctgctcgc ctgtgttgcc acctggattc tgcgcgggac gtccttctgc tacgtccctt
3301 cggccctcaa tccagcggac cttccttccc gcggcctgct gccggctctg cggcctcttc
3361 cgcgtcttcg ccttcgccct cagacgagtc ggatctccct ttgggccgcc tccccgcgtc
3421 gactttaaga ccaatgactt acaaggcagc tgtagatctt agccactttt taaaagaaaa
3481 gggggactg gaagggctaa ttcactccca acgaagacaa gatctgcttt ttgcttgtac
3541 tgggtctctc tggttagacc agatctgagc ctgggagctc tctggctaac tagggaaccc
3601 actgcttaag cctcaataaa gcttgccttg agtgcttcaa gtagtgtgtg cccgtctgtt
3661 gtgtgactct ggtaactaga gatccctcag acccttttag tcagtgtgga aaatctctag
3721 cagtacgtat agtagttcat gtcatcttat tattcagtat ttataacttg caaagaaatg
3781 aatatcagag agtgagagga acttgtttat tgcagcttat aatggttaca aataaagcaa
3841 tagcatcaca aatttcacaa ataaagcatt tttttcactg cattctagtt gtggtttgtc
3901 caaactcatc aatgtatctt atcatgtctg gctctagcta tcccgcccct aactccgccc
3961 atcccgcccc taactccgcc cagttccgcc cattctccgc cccatggctg actaattttt
4021 tttatttatg cagaggccga ggccgcctcg gcctctgagc tattccagaa gtagtgagga
4081 ggcttttttg gaggcctagg gacgtaccca attcgcccta tagtgagtcg tattacgcgc
4141 gctcactggc cgtcgtttta caacgtcgtg actgggaaaa ccctggcgtt acccaactta
4201 atcgccttgc agcacatccc cctttcgcca gctggcgtaa tagcgaagag gcccgcaccg
4261 atcgcccttc ccaacagttg cgcagcctga atggcgaatg ggacgcgccc tgtagcggcg
4321 cattaagcgc ggcgggtgtg gtggttacgc gcagcgtgac cgctacactt gccagcgccc
4381 tagcgcccgc tcctttcgct ttcttccctt cctttctcgc cacgttcgcc ggctttcccc
4441 gtcaagctct aaatcggggg ctccctttag ggttccgatt tagtgcttta cggcacctcg
4501 accccaaaaa acttgattag ggtgatggtt cacgtagtgg gccatcgccc tgatagacgg
4561 tttttcgccc tttgacgttg gagtccacgt tctttaatag tggactcttg ttccaaactg
4621 gaacaacact caaccctatc tcggtctatt cttttgattt ataagggatt ttgccgattt
4681 cggcctattg gttaaaaaat gagctgattt aacaaaaatt taacgcgaat tttaacaaaa
4741 tattaacgct tacaatttag gtggcacttt tcggggaaat gtgcgcggaa cccctatttg
4801 tttatttttc taaatacatt caaatatgta tccgctcatg agacaataac cctgataaat
4861 gcttcaataa tattgaaaaa ggaagagtat gagtattcaa catttccgtg tcgcccttat
4921 tccctttttt gcggcatttt gccttcctgt ttttgctcac ccagaaacgc tggtgaaagt
4981 aaaagatgct gaagatcagt tgggtgcacg agtgggttac atcgaactgg atctcaacag
5041 cggtaagatc cttgagagtt ttcgccccga agaacgtttt ccaatgatga gcacttttaa

5101 agttctgcta tgtggcgcgg tattatcccg tattgacgcc gggcaagagc aactcggtcg
5161 ccgcatacac tattctcaga atgacttggt tgagtactca ccagtcacag aaaagcatct
5221 tacggatggc atgacagtaa gagaattatg cagtgctgcc ataaccatga gtgataacac
5281 tgcggccaac ttacttctga caacgatcgg aggaccgaag gagctaaccg cttttttgca
5341 caacatgggg gatcatgtaa ctcgccttga tcgttgggaa ccggagctga atgaagccat
5401 accaaacgac gagcgtgaca ccacgatgcc tgtagcaatg gcaacaacgt tgcgcaaact
5461 attaactggc gaactactta ctctagcttc ccggcaacaa ttaatagact ggatggaggc
5521 ggataaagtt gcaggaccac ttctgcgctc ggcccttccg gctggctggt ttattgctga
5581 taaatctgga gccggtgagc gtgggtctcg cggtatcatt gcagcactgg ggccagatgg
5641 taagccctcc cgtatcgtag ttatctacac gacggggagt caggcaacta tggatgaacg
5701 aaatagacag atcgctgaga taggtgcctc actgattaag cattggtaac tgtcagacca
5761 agtttactca tatatacttt agattgattt aaaacttcat ttttaattta aaaggatcta
5821 ggtgaagatc ctttttgata atctcatgac caaaatccct taacgtgagt tttcgttcca
5881 ctgagcgtca gaccccgtag aaaagatcaa aggatcttct tgagatcctt tttttctgcg
5941 cgtaatctgc tgcttgcaaa caaaaaaacc accgctacca gcggtggttt gtttgccgga
6001 tcaagagcta ccaactcttt ttccgaaggt aactggcttc agcagagcgc agataccaaa
6061 tactgttctt ctagtgtagc cgtagttagg ccaccacttc aagaactctg tagcaccgcc
6121 tacatacctc gctctgctaa tcctgttacc agtggctgct gccagtggcg ataagtcgtg
6181 tcttaccggg ttggactcaa gacgatagtt accggataag gcgcagcggt cgggctgaac
6241 ggggggttcg tgcacacagc ccagcttgga gcgaacgacc tacaccgaac tgagatacct
6301 acagcgtgag ctatgagaaa gcgccacgct tcccgaaggg agaaaggcgg acaggtatcc
6361 ggtaagcggc agggtcggaa caggagagcg cacgagggag cttccagggg gaaacgcctg
6421 gtatctttat agtcctgtcg ggtttcgcca cctctgactt gagcgtcgat ttttgtgatg
6481 ctcgtcaggg gggcggagcc tatggaaaaa cgccagcaac gcggcctttt tacggttcct
6541 ggccttttgc tggccttttg ctcacatgtt ctttcctgcg ttatcccctg attctgtgga
6601 taaccgtatt accgcctttg agtgagctga taccgctcgc cgcagccgaa cgaccgagcg
6661 cagcgagtca gtgagcgagg aagcggaaga gcgcccaata cgcaaaccgc ctctccccgc
6721 gcgttggccg attcattaat gcagctggca cgacaggttt cccgactgga aagcgggcag
6781 tgagcgcaac gcaattaatg tgagttagct cactcattag gcaccccagg ctttacactt
6841 tatgcttccg gctcgtatgt tgtgtggaat tgtgagcgga taacaatttc acacaggaaa
6901 cagctatgac catgattacg ccaagcgcgc aattaaccct cactaaaggg aacaaaagct
6961 ggagctgcaa gcttaatgta gtcttatgca atactcttgt agtcttgcaa catggtaacg
7021 atgagttagc aacatgcctt acaaggagag aaaaagcacc gtgcatgccg attggtggaa
7081 gtaaggtggt acgatcgtgc cttattagga aggcaacaga cgggtctgac atggattgga
7141 cgaaccactg aattgccgca ttgcagagat attgtattta agtgcctagc tcgatacata
7201 aacgggtctc tctggttaga ccagatctga gcctgggagc tctctggcta actagggaac
7261 ccactgctta agcctcaata aagcttgcct tgagtgcttc aagtagtgtg tgcccgtctg
7321 ttgtgtgact ctggtaacta gagatccctc agaccctttt agtcagtgtg gaaaatctct
7381 agcagtggcg cccgaacagg gacttgaaag cgaaagggaa accagaggag ctctctcgac
7441 gcaggactcg gcttgctgaa gcgcgcacgg caagaggcga ggggcggcga ctggtgagta
7501 cgccaaaaat tttgactagc ggaggctaga aggagagaga tgggtgcgag agcgtcagta
7561 ttaagcgggg gagaattaga tcgcgatggg aaaaaattcg gttaaggcca gggggaaaga
7621 aaaaatataa attaaaacat atagtatggg caagcaggga gctagaacga ttcgcagtta
7681 atcctggcct gttagaaaca tcagaaggct gtagacaaat actgggacag ctacaaccat
7741 cccttcagac aggatcagaa gaacttagat cattatataa tacagtagca accctctatt
7801 gtgtgcatca aaggatagag ataaaagaca ccaaggaagc tttagacaag atagaggaag
7861 agcaaaacaa aagtaagacc accgcacagc aagcggccgc tgatcttcag acctggagga
7921 ggagatatga gggacaattg gagaagtgaa ttatataaat ataaagtagt aaaaattgaa
7981 ccattaggag tagcacccac caaggcaaag agaagagtgg tgcagagaga aaaaagagca
8041 gtgggaatag gagctttgtt ccttgggttc ttgggagcag caggaagcac tatgggcgca
8101 gcgtcaatga cgctgacggt acaggccaga caattattgt ctggtatagt gcagcagcag
8161 aacaatttgc tgagggctat tgaggcgcaa cagcatctgt tgcaactcac agtctggggc
8221 atcaagcagc tccaggcaag aatcctggct gtggaaagat acctaaagga tcaacagctc
8281 ctggggattt ggggttgctc tggaaaactc atttgcacca ctgctgtgcc ttggaatgct
8341 agttggagta ataaatctct ggaacagatt tggaatcaca cgacctggat ggagtgggac
8401 agagaaatta acaattacac aagcttaata cactccttaa ttgaagaatc gcaaaaccag
8461 caagaaaaga atgaacaaga attattggaa ttagataaat gggcaagttt gtggaattgg
8521 tttaacataa caaattggct gtggtatata aaattattca taatgatagt aggaggcttg
8581 gtaggtttaa gaatagtttt tgctgtactt tctatagtga atagagttag gcagggatat

```
8641 tcaccattat cgtttcagac ccacctccca accccgaggg gacccttgcg ccttttccaa
8701 ggcagccctg ggtttgcgca gggacgcggc tgctctgggc gtggttccgg gaaacgcagc
8761 ggcgccgacc ctgggtctcg cacattcttc acgtccgttc gcagcgtcac ccggatcttc
8821 gccgctaccc ttgtgggccc cccggcgacg cttcctgctc cgcccctaag tcgggaaggt
8881 tccttgcggt tcgcggcgtg ccggacgtga caaacggaag ccgcacgtct cactagtacc
8941 ctcgcagacg gacagcgcca gggagcaatg gcagcgcgcc gaccgcgatg ggctgtggcc
9001 aatagcggct gctcagcagg gcgcgccgag agcagcggcc gggaaggggc ggtgcgggag
9061 gcggggtgtg gggcggtagt gtgggccctg ttcctgcccg cgcggtgttc cgcattctgc
9121 aagcctccgg agcgcacgtc ggcagtcggc tccctcgttg accgaatcac cgacctctct
9181 ccccaggggg taccaccatg gccaagcctt tgtctcaaga agaatccacc ctcattgaaa
9241 gagcaacggc tacaatcaac agcatcccca tctctgaaga ctacagcgtc gccagcgcag
9301 ctctctctag cgacggccgc atcttcactg gtgtcaatgt atatcatttt actgggggac
9361 cttgtgcaga actcgtggtg ctgggcactg ctgctgctgc ggcagctggc aacctgactt
9421 gtatcgtcgc gatcggaaat gagaacaggg gcatcttgag cccctgcgga cggtgccgac
9481 aggtgcttct cgatctgcat cctgggatca aagccatagt gaaggacagt gatggacagc
9541 cgacggcagt tgggattcgt gaattgctgc cctctggtta tgtgtgggag ggcctgcagc
9601 tgcagtagta agaattctag atcttgagac aaatggcagt attcatccac aattttaaaa
9661 gaaaaggggg gattgggggg tacagtgcag gggaaagaat agtagacata atagcaacag
9721 acatacaaac taaagaatta caaaaacaaa ttacaaaaat tcaaaatttt cgggtttatt
9781 acagggacag cagagatcca ctttggcgcc ggctcgaggg g
//
```

pLX304-DCK*-IKZF1-IRES-GFP

Bicistronic lentiviral vector with IRES (internal ribosomal binding site) enabling CMV controlled expression of DCK*-IKZF1 (isoform Ik7) fusion protein, and eGFP (enhanced GFP).

DCK* encompasses three mutations in comparison to native *h.s.* deoxycytidine kinase (UniProt ID P27707): S74E; R104M; D133A.

Expressed fusion protein: DCK*-IKZF1-V5

```
1         10        20        30        40        50
|         |         |         |         |         |
MVPRGSHMATPPKRSCPSFSASSEGTRIKKISIEGNIAAGKSTFVNILKQ
LCEDWEVVPEPVARWCNVQSTQDEFEELTMEQKNGGNVLQMMYEKPERWS
FTFQTYACLSMIRAQLASLNGKLKDAEKPVLFFERSVYSARYIFASNLYE
SECMNETEWTIYQDWHDWMNNQFGQSLELDGIIYLQATPETCLHRIYLRG
RNEEQGIPLEYLEKLHYKHESWLLHRTLKTNFDYLQEVPILTLDVNEDFK
DKYESLVEKVKEFLSTLGGGSGGGSGGGSGGGSGGGSGGGSLSGSTSLYK
KVGMDADEGQDMSQVSGKESPPVSDTPDEGDEPMPIPEDLSTTSGGQQSS
KSDRVVASNVKVETQSDEENGRACEMNGEECAEDLRMLDASGEKMNGSHR
DQGSSALSGVGGIRLPNGKLKCDICGIICIGPNVLMVHKRSHTGERPFQC
NQCGASFTQKGNLLRHIKLHSGEKPFKCHLCNYACRRRDALTGHLRTHSV
IKEETNHSEMAEDLCKIGSERSLVLDRLASNVAKRKSSMPQKFLGDKGLS
DTPYDSSASYEKENEMMKSHVMDQAINNAINYLGAESLRPLVQTPPGGSE
VVPVISPMYQLHKPLAEGTPRSNHSAQDSAVENLLLLSKAKLVPSEREAS
PSNSCQDSTDTESNNEEQRSGLIYLTNHIAPHARNGLSLKEEHRAYDLLR
AASENSQDALRVVSTSGEQMKVYKCEHCRVLFLDHVMYTIHMGCHGFRDP
FECNMCGYHSQDRYEFSSHITRGEHRFHMSNPAFLYKVVGKPIPNPLLGL
DST
```

Expressed fluorescent marker: eGFP (enhanced GFP):

```
1         10        20        30        40        50
|         |         |         |         |         |
MVSKGEELFTGVVPILVELDGDVNGHKFSVSGEGEGDATYGKLTLKFICT
TGKLPVPWPTLVTTLTYGVQCFSRYPDHMKQHDFFKSAMPEGYVQERTIF
FKDDGNYKTRAEVKFEGDTLVNRIELKGIDFKEDGNILGHKLEYNYNSHN
VYIMADKQKNGIKVNFKIRHNIEDGSVQLADHYQQNTPIGDGPVLLPDNH
YLSTQSALSKDPNEKRDHMVLLEFVTAAGITLGMDELYK
```

Full plasmid sequence:

```
  1 gcccggggtt attaatagta atcaattacg gggtcattag ttcatagccc atatatggag
 61 ttccgcgtta cataacttac ggtaaatggc ccgcctggct gaccgcccaa cgacccccgc
121 ccattgacgt caataatgac gtatgttccc atagtaacgc caatagggac tttccattga
181 cgtcaatggg tggagtattt acggtaaact gcccacttgg cagtacatca agtgtatcat
241 atgccaagta cgccccctat tgacgtcaat gacggtaaat ggcccgcctg gcattatgcc
301 cagtacatga ccttatggga ctttcctact tggcagtaca tctacgtatt agtcatcgct
361 attaccatgg tgatgcggtt ttggcagtac atcaatgggc gtggatagcg gtttgactca
421 cggggatttc caagtctcca ccccattgac gtcaatggga gtttgttttg gcaccaaaat
481 caacgggact ttccaaaatg tcgtaacaac tccgccccat tgacgcaaat gggcggtagg
541 cgtgtacggt gggaggtcta tataagcaga gctctctggc taagccacca tggttccgcg
601 tggctctcat atggccaccc cgcccaagag aagctgccca tctttctcag ccagctctga
661 ggggacccgc atcaagaaaa tctccatcga agggaacatc gctgcaggga agtcaacatt
721 tgtgaatatc cttaaacaat tgtgtgaaga ttgggaagtg gttcctgaac ctgttgccag
781 atggtgcaat gttcaaagta ctcaagatga atttgaggaa cttacaatgg agcagaaaaa
```

841 tggtgggaat gttcttcaga tgatgtatga gaaacctgaa cgatggtctt ttaccttcca
901 aacctacgcc tgtctcagta tgataagagc tcagcttgcc tctctgaatg gcaagctcaa
961 agatgcagag aaacctgtat tattttttga acgatctgtg tatagtgcga ggtatatttt
1021 tgcatctaat ttgtatgaat ctgaatgcat gaatgagaca gagtggacaa tttatcaaga
1081 ctggcatgac tggatgaata accaatttgg ccaaagcctt gaattggatg gaatcattta
1141 tcttcaagcc actccagaga catgcttaca tagaatatat ttacggggaa gaaatgaaga
1201 gcaaggcatt cctcttgaat atttagagaa gcttcattat aaacatgaaa gctggctcct
1261 gcataggaca ctgaaaacca acttcgatta tcttcaagag gtgcctatct taacactgga
1321 tgttaatgaa gactttaaag acaaatatga aagtctggtt gaaaaggtca aagagttttt
1381 gagtactttg ggagggggta gcggcggagg ttcaggaggc ggaagtggtg gtggctccgg
1441 aggcggtagt ggcggaggtt cactgtcggg atcaacaagt ttgtacaaaa aagttggcat
1501 ggatgctgat gagggtcaag acatgtccca agtttcaggg aaggaaagcc cccctgtaag
1561 cgatactcca gatgagggcg atgagcccat gccgatcccc gaggacctct ccaccacctc
1621 gggaggacag caaagctcca agagtgacag agtcgtggcc agtaatgtta aagtagagac
1681 tcagagtgat gaagagaatg ggcgtgcctg tgaaatgaat ggggaagaat gtgcggagga
1741 tttacgaatg cttgatgcct cgggagagaa aatgaatggc tcccacaggg accaaggcag
1801 ctcggctttg tcgggagttg gaggcattcg acttcctaac ggaaaactaa agtgtgatat
1861 ctgtgggatc atttgcatcg ggcccaatgt gctcatggtt cacaaaagaa gccacactgg
1921 agaacggccc ttccagtgca atcagtgcgg ggcctcattc acccagaagg gcaacctgct
1981 ccggcacatc aagctgcatt ccggggagaa gcccttcaaa tgccacctct gcaactacgc
2041 ctgccgccgg agggacgccc tcactggcca cctgaggacg cactccgtca ttaaagaaga
2101 aactaatcac agtgaaatgg cagaagacct gtgcaagata ggatcagaga gatctctcgt
2161 gctggacaga ctagcaagta acgtcgccaa acgtaagagc tctatgcctc agaaatttct
2221 tggggacaag ggcctgtccg acacgcccta cgacagcagc gccagctacg agaaggagaa
2281 cgaaatgatg aagtcccacg tgatggacca agccatcaac aacgccatca actacctggg
2341 ggccgagtcc ctgcgcccgc tggtgcagac gcccccgggc ggttccgagg tggtcccggt
2401 catcagcccg atgtaccagc tgcacaagcc gctcgcggag ggcaccccgc gctccaacca
2461 ctcggcccag gacagcgccg tggagaacct gctgctgctc tccaaggcca agttggtgcc
2521 ctcggagcgc gaggcgtccc cgagcaacag ctgccaagac tccacggaca ccgagagcaa
2581 caacgaggag cagcgcagcg gtctcatcta cctgaccaac cacatcgccc cgcacgcgcg
2641 caacgggctg tcgctcaagg aggagcaccg cgcctacgac ctgctgcgcg ccgcctccga
2701 gaactcgcag gacgcgctcc gcgtggtcag caccagcggg gagcagatga aggtgtacaa
2761 gtgcgaacac tgccgggtgc tcttcctgga tcacgtcatg tacaccatcc acatgggctg
2821 ccacggcttc cgtgatcctt ttgagtgcaa catgtgcggc taccacagcc aggaccggta
2881 cgagttctcg tcgcacataa cgcgagggga gcaccgcttc cacatgagca acccagcttt
2941 cttgtacaaa gtggttggta agcctatccc taaccctctc ctcggtctcg attctacgta
3001 gtaatgagct agccgctacg taaattccgc cccccccccc cctctccctc cccccccccct
3061 aacgttactg gccgaagccg cttggaataa ggccggtgtg cgtttgtcta tatgttattt
3121 tccaccatat tgccgtcttt tggcaatgtg agggcccgga aacctggccc tgtcttcttg
3181 acgagcattc ctaggggtct ttcccctctc gccaaaggaa tgcaaggtct gttgaatgtc
3241 gtgaaggaag cagttcctct ggaagcttct tgaagacaaa caacgtctgt agcgaccctt
3301 tgcaggcagc ggaacccccc acctggcgac aggtgcctct gcggccaaaa gccacgtgta
3361 taagatacac ctgcaaaggc ggcacaaccc cagtgccacg ttgtgagttg gatagttgtg
3421 gaaagagtca aatggctctc ctcaagcgta ttcaacaagg ggctgaagga tgcccagaag
3481 gtaccccatt gtatgggatc tgatctgggg cctcggtgca catgctttac atgtgtttag
3541 tcgaggttaa aaaaacgtct aggccccccg aaccacgggg acgtggtttt cctttgaaaa
3601 acacgatgat aatatggcca caaccatggt gagcaagggc gaggagctgt tcaccggggt
3661 ggtgcccatc ctggtcgagc tggacggcga cgtaaacggc cacaagttca gcgtgtccgg
3721 cgagggcgag ggcgatgcca cctacggcaa gctgaccctg aagttcatct gcaccaccgg
3781 caagctgccc gtgccctggc ccaccctcgt gaccaccctg acctacggcg tgcagtgctt
3841 cagccgctac cccgaccaca tgaagcagca cgacttcttc aagtccgcca tgcccgaagg
3901 ctacgtccag gagcgcacca tcttcttcaa ggacgacggc aactacaaga cccgcgccga
3961 ggtgaagttc gagggcgaca ccctggtgaa ccgcatcgag ctgaagggca tcgacttcaa
4021 ggaggacggc aacatcctgg ggcacaagct ggagtacaac tacaacagcc acaacgtcta
4081 tatcatggcc gacaagcaga agaacggcat caaggtgaac ttcaagatcc gccacaacat
4141 cgaggacggc agcgtgcagc tcgccgacca ctaccagcag aacaccccca tcggcgacgg
4201 ccccgtgctg ctgcccgaca accactacct gagcacccag tccgccctga gcaaagaccc
4261 caacgagaag cgcgatcaca tggtcctgct ggagttcgtg accgccgccg ggatcactct
4321 cggcatggac gagctgtaca agtaaaccgg tggcgcgtta agtcgacaat caacctctgg

4381 attacaaaat ttgtgaaaga ttgactggta ttcttaacta tgttgctcct tttacgctat
4441 gtggatacgc tgctttaatg cctttgtatc atgctattgc ttcccgtatg gctttcattt
4501 tctcctcctt gtataaatcc tggttgctgt ctctttatga ggagttgtgg cccgttgtca
4561 ggcaacgtgg cgtggtgtgc actgtgtttg ctgacgcaac ccccactggt tggggcattg
4621 ccaccacctg tcagctcctt tccgggactt tcgctttccc cctccctatt gccacggcgg
4681 aactcatcgc cgcctgcctt gcccgctgct ggacaggggc tcggctgttg ggcactgaca
4741 attccgtggt gttgtcgggg aaatcatcgt cctttccttg gctgctcgcc tgtgttgcca
4801 cctggattct gcgcgggacg tccttctgct acgtcccttc ggccctcaat ccagcggacc
4861 ttccttcccg cggcctgctg ccggctctgc ggcctcttcc gcgtcttcgc cttcgccctc
4921 agacgagtcg gatctccctt tgggccgcct ccccgcgtcg actttaagac caatgactta
4981 caaggcagct gtagatctta gccacttttt aaaagaaaag gggggactgg aagggctaat
5041 tcactcccaa cgaagacaag atctgctttt tgcttgtact gggtctctct ggttagacca
5101 gatctgagcc tgggagctct ctggctaact agggaaccca ctgcttaagc ctcaataaag
5161 cttgccttga gtgcttcaag tagtgtgtgc ccgtctgttg tgtgactctg gtaactagag
5221 atccctcaga cccttttagt cagtgtggaa aatctctagc agtacgtata gtagttcatg
5281 tcatcttatt attcagtatt tataacttgc aaagaaatga atatcagaga gtgagaggaa
5341 cttgtttatt gcagcttata atggttacaa ataaagcaat agcatcacaa atttcacaaa
5401 taaagcattt ttttcactgc attctagttg tggtttgtcc aaactcatca atgtatctta
5461 tcatgtctgg ctctagctat cccgccccta actccgccca tcccgcccct aactccgccc
5521 agttccgccc attctccgcc ccatggctga ctaatttttt ttatttatgc agaggccgag
5581 gccgcctcgg cctctgagct attccagaag tagtgaggag gcttttttgg aggcctaggg
5641 acgtacccaa ttcgccctat agtgagtcgt attacgcgcg ctcactggcc gtcgttttac
5701 aacgtcgtga ctgggaaaac cctggcgtta cccaacttaa tcgccttgca gcacatcccc
5761 ctttcgccag ctggcgtaat agcgaagagg cccgcaccga tcgcccttcc caacagttgc
5821 gcagcctgaa tggcgaatgg gacgcgccct gtagcggcgc attaagcgcg gcgggtgtgg
5881 tggttacgcg cagcgtgacc gctacacttg ccagcgccct agcgcccgct cctttcgctt
5941 tcttcccttc ctttctcgcc acgttcgccg gctttccccg tcaagctcta aatcgggggc
6001 tccctttagg gttccgattt agtgctttac ggcacctcga ccccaaaaaa cttgattagg
6061 gtgatggttc acgtagtggg ccatcgccct gatagacggt ttttcgccct ttgacgttgg
6121 agtccacgtt ctttaatagt ggactcttgt tccaaactgg aacaacactc aaccctatct
6181 cggtctattc ttttgattta taagggattt tgccgatttc ggcctattgg ttaaaaaatg
6241 agctgattta acaaaaattt aacgcgaatt ttaacaaaat attaacgctt acaatttagg
6301 tggcactttt cggggaaatg tgcgcggaac ccctatttgt ttatttttct aaatacattc
6361 aaatatgtat ccgctcatga gacaataacc ctgataaatg cttcaataat attgaaaaag
6421 gaagagtatg agtattcaac atttccgtgt cgcccttatt cccttttttg cggcattttg
6481 ccttcctgtt tttgctcacc cagaaacgct ggtgaaagta aaagatgctg aagatcagtt
6541 gggtgcacga gtgggttaca tcgaactgga tctcaacagc ggtaagatcc ttgagagttt
6601 tcgccccgaa gaacgttttc caatgatgag cacttttaaa gttctgctat gtggcgcggt
6661 attatcccgt attgacgccg ggcaagagca actcggtcgc cgcatacact attctcagaa
6721 tgacttggtt gagtactcac cagtcacaga aaagcatctt acggatggca tgacagtaag
6781 agaattatgc agtgctgcca taaccatgag tgataacact gcggccaact tacttctgac
6841 aacgatcgga ggaccgaagg agctaaccgc ttttttgcac aacatggggg atcatgtaac
6901 tcgccttgat cgttgggaac cggagctgaa tgaagccata ccaaacgacg agcgtgacac
6961 cacgatgcct gtagcaatgg caacaacgtt gcgcaaacta ttaactggcg aactacttac
7021 tctagcttcc cggcaacaat taatagactg gatggaggcg gataaagttg caggaccact
7081 tctgcgctcg gcccttccgg ctggctggtt tattgctgat aaatctggag ccggtgagcg
7141 tgggtctcgc ggtatcattg cagcactggg gccagatggt aagccctccc gtatcgtagt
7201 tatctacacg acggggagtc aggcaactat ggatgaacga aatagacaga tcgctgagat
7261 aggtgcctca ctgattaagc attggtaact gtcagaccaa gtttactcat atatacttta
7321 gattgattta aaacttcatt tttaatttaa aaggatctag gtgaagatcc tttttgataa
7381 tctcatgacc aaaatccctt aacgtgagtt ttcgttccac tgagcgtcag accccgtaga
7441 aaagatcaaa ggatcttctt gagatccttt ttttctgcgc gtaatctgct gcttgcaaac
7501 aaaaaaacca ccgctaccag cggtggtttg tttgccggat caagagctac caactctttt
7561 tccgaaggta actggcttca gcagagcgca gataccaaat actgttcttc tagtgtagcc
7621 gtagttaggc caccacttca agaactctgt agcaccgcct acatacctcg ctctgctaat
7681 cctgttacca gtggctgctg ccagtggcga taagtcgtgt cttaccgggt tggactcaag
7741 acgatagtta ccggataagg cgcagcggtc gggctgaacg gggggttcgt gcacacagcc
7801 cagcttggag cgaacgacct acaccgaact gagataccta cagcgtgagc tatgagaaag
7861 cgccacgctt cccgaaggga gaaaggcgga caggtatccg gtaagcggca gggtcggaac

7921 aggagagcgc acgagggagc ttccaggggg aaacgcctgg tatctttata gtcctgtcgg
7981 gtttcgccac ctctgacttg agcgtcgatt tttgtgatgc tcgtcagggg ggcggagcct
8041 atggaaaaac gccagcaacg cggccttttt acggttcctg gccttttgct ggccttttgc
8101 tcacatgttc tttcctgcgt tatcccctga ttctgtggat aaccgtatta ccgcctttga
8161 gtgagctgat accgctcgcc gcagccgaac gaccgagcgc agcgagtcag tgagcgagga
8221 agcggaagag cgcccaatac gcaaaccgcc tctccccgcg cgttggccga ttcattaatg
8281 cagctggcac gacaggtttc ccgactggaa agcgggcagt gagcgcaacg caattaatgt
8341 gagttagctc actcattagg caccccaggc tttacacttt atgcttccgg ctcgtatgtt
8401 gtgtggaatt gtgagcggat aacaatttca cacaggaaac agctatgacc atgattacgc
8461 caagcgcgca attaaccctc actaaaggga acaaaagctg gagctgcaag cttaatgtag
8521 tcttatgcaa tactcttgta gtcttgcaac atggtaacga tgagttagca acatgcctta
8581 caaggagaga aaaagcaccg tgcatgccga ttggtggaag taaggtggta cgatcgtgcc
8641 ttattaggaa ggcaacagac gggtctgaca tggattggac gaaccactga attgccgcat
8701 tgcagagata ttgtatttaa gtgcctagct cgatacataa acgggtctct ctggttagac
8761 cagatctgag cctgggagct ctctggctaa ctagggaacc cactgcttaa gcctcaataa
8821 agcttgcctt gagtgcttca agtagtgtgt gcccgtctgt tgtgtgactc tggtaactag
8881 agatccctca gaccctttta gtcagtgtgg aaaatctcta gcagtggcgc ccgaacaggg
8941 acttgaaagc gaaagggaaa ccagaggagc tctctcgacg caggactcgg cttgctgaag
9001 cgcgcacggc aagaggcgag gggcggcgac tggtgagtac gccaaaaatt ttgactagcg
9061 gaggctagaa ggagagagat gggtgcgaga gcgtcagtat taagcggggg agaattagat
9121 cgcgatggga aaaaattcgg ttaaggccag ggggaaagaa aaaatataaa ttaaaacata
9181 tagtatgggc aagcagggag ctagaacgat tcgcagttaa tcctggcctg ttagaaacat
9241 cagaaggctg tagacaaata ctgggacagc tacaaccatc ccttcagaca ggatcagaag
9301 aacttagatc attatataat acagtagcaa ccctctattg tgtgcatcaa aggatagaga
9361 taaaagacac caaggaagct ttagacaaga tagaggaaga gcaaaacaaa agtaagacca
9421 ccgcacagca agcggccgct gatcttcaga cctggaggag gagatatgag ggacaattgg
9481 agaagtgaat tatataaata taaagtagta aaaattgaac cattaggagt agcacccacc
9541 aaggcaaaga gaagagtggt gcagagagaa aaaagagcag tgggaatagg agctttgttc
9601 cttgggttct tgggagcagc aggaagcact atgggcgcag cgtcaatgac gctgacggta
9661 caggccagac aattattgtc tggtatagtg cagcagcaga acaatttgct gagggctatt
9721 gaggcgcaac agcatctgtt gcaactcaca gtctggggca tcaagcagct ccaggcaaga
9781 atcctggctg tggaaagata cctaaaggat caacagctcc tggggatttg gggttgctct
9841 ggaaaactca tttgcaccac tgctgtgcct tggaatgcta gttggagtaa taaatctctg
9901 gaacagattt ggaatcacac gacctggatg gagtgggaca gagaaattaa caattacaca
9961 agcttaatac actccttaat tgaagaatcg caaaaccagc aagaaaagaa tgaacaagaa
10021 ttattggaat tagataaatg ggcaagtttg tggaattggt ttaacataac aaattggctg
10081 tggtatataa aattattcat aatgatagta ggaggcttgg taggtttaag aatagttttt
10141 gctgtacttt ctatagtgaa tagagttagg cagggatatt caccattatc gtttcagacc
10201 cacctcccaa ccccgagggg acccttgcgc cttttccaag gcagccctgg gtttgcgcag
10261 ggacgcggct gctctgggcg tggttccggg aaacgcagcg gcgccgaccc tgggtctcgc
10321 acattcttca cgtccgttcg cagcgtcacc cggatcttcg ccgctaccct tgtgggcccc
10381 ccggcgacgc ttcctgctcc gccccctaagt cgggaaggtt ccttgcggtt cgcggcgtgc
10441 cggacgtgac aaacggaagc cgcacgtctc actagtaccc tcgcagacgg acagcgccag
10501 ggagcaatgg cagcgcgccg accgcgatgg gctgtggcca atagcggctg ctcagcaggg
10561 cgcgccgaga gcagcggccg ggaaggggcg gtgcgggagg cggggtgtgg ggcggtagtg
10621 tgggccctgt tcctgcccgc gcggtgttcc gcattctgca agcctccgga gcgcacgtcg
10681 gcagtcggct ccctcgttga ccgaatcacc gacctctctc cccagggggt accaccatgg
10741 ccaagccttt gtctcaagaa gaatccaccc tcattgaaag agcaacggct acaatcaaca
10801 gcatccccat ctctgaagac tacagcgtcg ccagcgcagc tctctctagc gacggccgca
10861 tcttcactgg tgtcaatgta tatcatttta ctgggggacc ttgtgcagaa ctcgtggtgc
10921 tgggcactgc tgctgctgcg gcagctggca acctgacttg tatcgtcgcg atcggaaatg
10981 agaacagggg catcttgagc ccctgcggac ggtgccgaca ggtgcttctc gatctgcatc
11041 ctgggatcaa agccatagtg aaggacagtg atggacagcc gacggcagtt gggattcgtg
11101 aattgctgcc ctctggttat gtgtgggagg gcctgcagct gcagtagtaa gaattctaga
11161 tcttgagaca aatggcagta ttcatccaca attttaaaag aaaagggggg attggggggt
11221 acagtgcagg ggaaagaata gtagacataa tagcaacaga catacaaact aaagaattac
11281 aaaaacaaat tacaaaaatt caaaattttc gggtttatta cagggacagc agagatccac
11341 tttggcgccg gctcgagggg
//
